# Reducing 14-3-3ζ expression influences adipocyte maturity and impairs function

**DOI:** 10.1101/2020.02.24.962662

**Authors:** Abel K. Oppong, Kadidia Diallo, Isabelle Robillard Frayne, Christine Des Rosiers, Gareth E. Lim

**Author notes:** To whom correspondence should be address: Gareth E. Lim, CRCHUM, Tour Viger, Rm 08.482, 900 Rue St. Denis, Montréal, QC H2X 029, Canada.

## Abstract

One of the primary metabolic functions of a mature adipocyte is to supply energy via lipolysis, or the catabolism of stored lipids. Hormone-sensitive lipase (HSL) is a critical lipolytic enzyme, and its phosphorylation and subsequent activation by PKA generates phospho-binding sites for 14-3-3 proteins, a ubiquitously expressed family of molecular scaffolds. While we previously identified essential roles of the 14-3-3ζ isoform in murine adipogenesis, the presence of 14-3-3 protein binding sites on HSL suggests that 14-3-3ζ could also influence mature adipocyte processes like lipolysis. Herein, we demonstrate that 14-3-3ζ is necessary for lipolysis in male mice and fully differentiated 3T3-L1 adipocytes, as depletion of 14-3-3ζ significantly impaired glycerol and FFA release. Unexpectedly, this was not due to impairments in signaling events underlying lipolysis; instead, reducing 14-3-3ζ expression was found to significantly impact adipocyte maturity, as observed by reduced abundance of PPARγ2 protein and expression of mature adipocytes genes and those associated with *de novo* triglyceride synthesis and lipolysis. The impact of 14-3-3ζ depletion on adipocyte maturity was further examined with untargeted lipidomics, which revealed that reductions in 14-3-3ζ abundance promoted the acquisition of a lipidomic signature that resembled undifferentiated, pre-adipocytes. Collectively, these findings reveal a novel aspect of 14-3-3ζ in adipocytes, as reducing 14-3-3ζ was found to have a negative effect on adipocyte maturity and adipocyte-specific processes like lipolysis.

## Introduction

The primary function of white adipose tissue (WAT) is to regulate energy and nutrient homeostasis, as white adipocytes specialize in both the storage of triacylglycerols (TAGs) and the mobilization of free fatty acids (FFAs). These processes occur in response to metabolic demands associated with prolonged periods of activity or fasting (17). In mature adipocytes, FFAs are generated by the hydrolysis of TAGs through a process known as lipolysis, which occurs following the binding of catecholamines to β3-adrenergic receptors, the subsequent generation of the second messenger, cAMP, and the activation of Protein Kinase A (PKA) (17). TAG hydrolysis is principally mediated by three lipases: adipose triacylglycerol lipase (ATGL), hormone-sensitive lipase (HSL), and monoacylglycerol lipase (MAGL), whereby they respectively catalyze the sequential conversion of TAGs into diacylglycerols (DAGs), monoacylglycerols (MAGs), and finally FFAs and glycerol (3, 23).

Molecular scaffolds belonging to the 14-3-3 family are ubiquitously expressed proteins that have been implicated in regulating cellular processes, such as proliferation, apoptosis and metabolism (19, 34, 41). These functions arise from their ability to interact with target proteins harboring specific phosphorylated serine and threonine motifs to modulate their activities or subcellular localization (16, 21). For example, the inhibitory action of the Rab-GAP, AS160/TBC1D4, on GLUT4 translocation is attenuated following its interaction with 14-3-3 proteins (45), and binding of FOXO1 to 14-3-3 proteins promotes its retention in the cytoplasm (6). We previously reported essential roles of one of the seven mammalian isoforms, 14-3-3ζ, in adipogenesis, as systemic deletion of 14-3-3ζ in mice resulted in significant reductions in visceral adipose tissue mass and decreased expression of mature adipocyte markers in gonadal adipocytes (34). Furthermore, through the use of proteomics to elucidate the 14-3-3ζ interactome, we found that 14-3-3ζ is necessary for RNA splicing during adipocyte differentiation (42). However, whether 14-3-3ζ can also influence metabolic processes specific to mature adipocytes has not been fully examined.

One of the earliest reported functions of 14-3-3 proteins is their regulatory functions on tyrosine and tryptophan hydroxylases, which accounts for their alternative names of tyrosine 3-monooxygenase/tryptophan 5-monooxygenase activation proteins (28). Given their ability to bind to proteins harboring specific serine or threonine motifs, they can regulate a diverse array of enzymes and influence intracellular signaling pathways. For example, the activity of other kinases, such as PKA and RAF-1, are positively regulated upon binding to 14-3-3 proteins, but in some instances, inhibitory effects, as seen with DYRK1A, have been reported (18, 25, 26). Phosphorylation of HSL by PKA generates 14-3-3 protein binding sites (36), and with the ability of 14-3-3 proteins to broadly regulate enzyme activity, we initially hypothesized that alterations in 14-3-3ζ expression in adipocytes could impact HSL function, thereby influencing lipolysis.

Herein, we report that reducing 14-3-3ζ protein abundance in mature adipocytes impaired lipolysis; however, this was due to changes in the maturity of adipocytes and not caused by defects associated with downstream signaling events in the β3-adrenergic signaling pathway. Signficant reductions in mRNA and protein levels for key markers of mature adipocytes, including *Pparg*/PPARγ, *Hsl*/HSL, and *Atgl*/ATGL were observed, in addition to decreased mRNA levels for genes involved in *de novo* triglyceride synthesis and fatty acid transport. To further explore the maturity of 14-3-3ζ-depleted adipocytes, untargeted lipidomics was performed, and reductions in 14-3-3ζ protein abundance was found to promote the acquisition of a lipid signature resembling pre-adipocytes. In conclusion, this study identifies critical roles of 14-3-3ζ in regulating adipocyte maturity, which can influence overall adipocyte processes like lipolysis.

## Materials and Methods

### Animal husbandry

*Adipoq*-CreERT2^Soff^ mice (stock no. 025124, JAX, Bar Harbor Maine) were bred with *Ywhaz*^flox/flox^ mice harboring LoxP sites that flank exon 4 of *Ywhaz* (gene that encodes 14-3-3ζ) (Ywhaz^tm1c(EUCOMM)HMgu^, Toronto Centre for Phenogenomics, Toronto, Canada) to generate *Adipoq-*CreERT2-14-3-3ζKO mice (adi14-3-3ζKO) (14). Both strains were on a C57Bl/6J background. To delete exon 4 of *Ywhaz*, tamoxifen (TMX, 50 mg/kg b.w; Sigma Aldrich, St. Louis, MO) was administered by intraperitoneal injections for five days (Figure 1A). Transgenic mice over-expressing a TAP-tagged human 14-3-3ζ molecule were on a CD1 background and were provided by the laboratory of Dr. Amparo Acker-Palmer (2, 34). All mice were maintained on a standard chow diet (Teklad diet no. TD2918) under 12-hour light/12-hour dark cycles in an environmentally controlled setting (22°C ± 1°C) with free access to food and water. All procedures were approved and performed in accordance with CIPA (Comité institutionnel de protection des animaux du CRCHUM) guidelines at the University of Montreal Hospital Research Centre.

**Figure 1.**
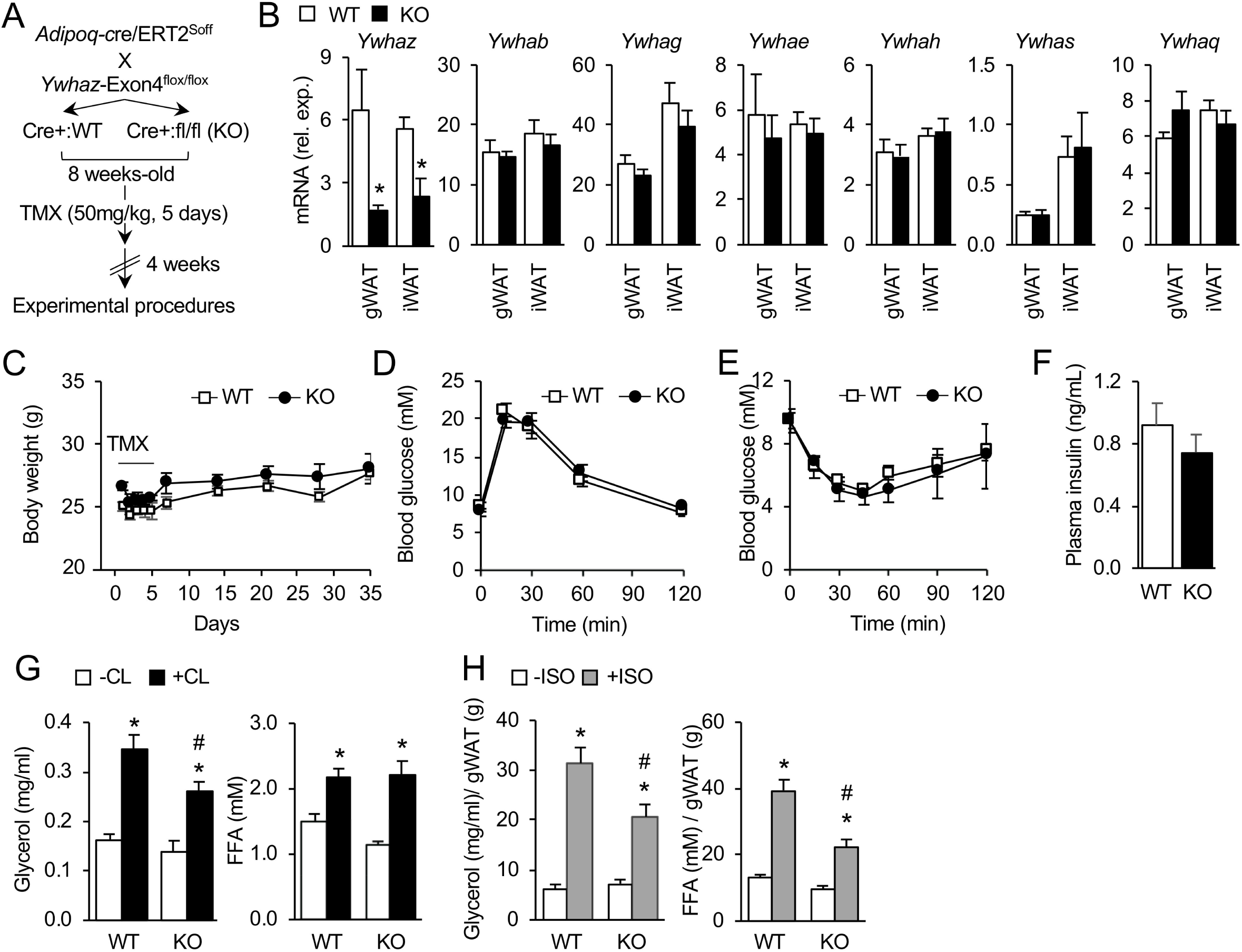
Deletion of 14-3-3ζ in adipocytes impairs lipolysis. **(A)** Experimental design to examine the role of 14-3-3ζ in lipolysis. *Adipoq-*CreERT2^Soff^ mice were bred with those harboring Loxp sites flanking exon4 of *Ywhaz*, the gene encoding 14-3-3ζ. At 8-weeks of age, wildtype (WT) or adi14-3-3ζKO (KO) mice were injected with tamoxifen (TMX, 50 mg/kg; 5 days), and mice were use for experimental procedures 4 weeks after the last TMX injection. **(B)** Levels of *Ywhaz, Ywhab, Ywhag, Ywhae, Ywhah, Ywhas* and *Ywhaq* (encoding 14-3-3ζ, 14-3-3β, 14-3-3γ, 14-3-3ε, 14-3-3η, 14-3-3σ, and 14-3-3θ, respectively) in gonadal (gWAT) and inguinal (iWAT) adipose depots of male WT or adi14-3-3ζKO mice (n=8-9 WT, n=5 KO for male mice; two-way ANOVA, followed by Bonferroni t-tests; *: p<0.05 when compared to WT). **(C)** Body weights of male adi14-3-3ζKO mice following i.p injections of tamoxifen (50 mg/kg b.w) were measured weekly (n=9 WT, n=6 KO for male mice). **(D**,**E)** Intraperitoneal glucose (D, 2 g/kg b.w.) and insulin (E, 0.5 U/kg b.w.) tolerance tests on male adi14-3-3ζKO mice fasted for 6 and 4 hours, respectively, at 12 weeks of age (n=9 WT, n=6 KO for male mice). **(F)** Plasma insulin levels from WT and adi14-3-3ζKO mice following an overnight fast (n=9 WT, n=6 KO**). (G)** Plasma glycerol and FFA levels in male adi14-3-3ζKO mice after injection with CL-316,243 (CL, 1 mg/kg b.w.), following an overnight fast (n=9 WT, n=6 KO; two-way ANOVA, followed by Bonferroni t-tests; *: p<0.05 when compared to -CL, #: p<0.05 when compared to WT). **(H)** Glycerol and FFA levels in the supernatant of gonadal adipose tissue explants treated with isoproterenol (Iso, 1 μM) for 2 hours. Glycerol and FFA release were normalized to the mass of the explants (n=5 WT, n=3 KO; two-way ANOVA, followed by Bonferroni t-tests; #: p<0.05 when compared to WT). All values represent mean ± SE.

### Metabolic phenotyping

For glucose tolerance tests, adi14-3-3ζKO mice were fasted for six hours and challenged with d-glucose (2 g/kg b.w.; VWR, Solon, OH) by intraperitoneal administration (34, 35). For insulin tolerance tests, adi14-3-3ζKO mice were fasted for four hours and were injected intraperitoneally with Humulin R insulin (0.5 U/kg b.w; Eli Lilly, Toronto, ON) (34, 35). Blood glucose was measured from tail blood with a Contour Next EZ glucose meter (Ascencia Diabetes Care, Basel, Switzerland). Plasma leptin, insulin, and adipocnectin were measured by ELISA (ALPCO, Keewatin, NH) according to the manufacturer’s protocols.

### Cell Culture and transient transfections

3T3-L1 cells were maintained in DMEM (Life Technologies Corporation, Grand Island, NY), supplemented with 10% newborn calf serum (NBCS) and 1% penicillin/streptomycin (P/S) and were seeded onto 12-well plates (100,000 cells/well) or 10cm dishes (2×10^6^ cells/dish) 2 days prior to the induction of differentiation. Confluent cells were treated with a differentiation cocktail (DMEM supplemented with 10% FBS, 1% P/S, 500 μM IBMX, 500 nM dexamethasone and 172 nM insulin) for 48 hours, followed by media replacement (DMEM with 10% FBS, 1% P/S and 172 nM insulin) every 2 days for 7-8 days (34, 42). Knockdown and overexpression of 14-3-3ζ was achieved by transfecting day 7-8 differentiated cells with scrambled control siRNA, 14-3-3ζ-specific siRNA (Ambion, Austin, TX), GFP control plasmids, or plasmids encoding 14-3-3ζ (14-3-3ζ IRES-GFP) using the Amaxa Cell Line Nucleofector Kit L, as per manufacturer’s instructions (Lonza, Koln, Germany). Differentiated 3T3-L1 cells were stained with Oil Red-O (ORO) and measured, as previously described (34, 42). All studies with 3T3-L1 cells were performed on cells between passages 10-16.

### Measurements of lipolysis

Adi14-3-3ζKO mice were injected with 1 mg/kg CL-316,243 (Sigma-Aldrich, St. Louis, MO) following an overnight fast. TAP mice were injected with 10 mg/kg isoproterenol (Sigma-Aldrich), following an overnight fast. Blood was collected from the tail vein before and 30 minutes after administration of CL-316,243 or isoproterenol. To measure lipolysis *ex vivo*, mice where sacrificed 4 weeks after the last TMX injection, and gonadal adipose tissues were harvested and placed into pH 7.4-adjusted Krebs-Ringer buffer (135 mM NaCl, 3.6 mM KCl, 0.5 mM NaH_2_PO_4_, 0.5 mM MgCl_2_, and 1.5 mM CaCl_2_), supplemented with 10 mM HEPES, 2 mM NaHCO_3_, 5 mM glucose and 2% bovine serum albumin (BSA). Gonadal explants were treated with or without 1 μM isoproterenol (Sigma Aldrich) for two hours. The supernatant was collected and centrifuged for 15 minutes at 1500 RPM. To measure lipolysis from differentiated 3T3-L1 cells (Zenbio, Research Triangle Park, NC), cells were incubated in starvation media consisting of pH 7.4-adjusted Krebs-Ringer buffer, 5 mM glucose and 0.2% BSA for two hours. Cells were then incubated in experimental media consisting of pH 7.4-adjusted Krebs-Ringer buffer, 5 mM glucose and 2% BSA for two hours while treated with 1 μM isoproterenol (Sigma-Aldrich), 10 μM forskolin with 0.5 mM IBMX (Sigma-Aldrich), or 1 mM dibutyryl cAMP (Sigma-Aldrich). Lipolysis was assessed by measuring glycerol and free fatty acid (FFA) levels using triglyceride (Sigma-Aldrich) and non-esterified fatty acid (NEFA; Wako Diagnostics, Osaka, Japan) kits, as per the manufacturer’s protocol.

### RNA isolation and qPCR

After 48 hours, total RNA was isolated from differentiated 3T3-L1 cells using the RNeasy kit (Qiagen, Hilden, Germany). cDNA was generated using the High-Capacity cDNA Reverse Transcription kit (ThermoFisher Scientific, Waltham, MA). Measurements of mRNA were performed with SYBR green chemistry using the QuantStudio 6-flex Real-time PCR System (ThermoFisher Scientific). All data were normalized to either *Hprt* or *Actinb* by the ΔC(t) method (34, 42). Primer sequences are listed in Supplemental Table 1.

### Measurement of intracellular cAMP levels

After 7 days of differentiation, mature 3T3-L1 adipocytes were transfected with scrambled, control siRNA (siCon) or siRNA against 14-3-3ζ (si14-3-3ζ). After 48 hours, cells were incubated in starvation media (pH 7.4-adjusted Krebs-Ringer buffer, 5 mM glucose, and 0.2% BSA) for two hours, followed by incubation in experimental media in the presence of 1 μM Isoproterenol or 20 μM forskolin with 0.5 mM IBMX for one hour. Lysates were harvested and intracellular cAMP levels were assayed using the cAMP Parameter Assay Kit (R&D Systems, Minneapolis, MN).

### Immunoblotting and immunohistochemistry

Differentiated 3T3-L1 cells were lysed in RIPA buffer (0.9% NaCl, 1% v/v triton X-100, 0.5% sodium deoxycholate, 0.1% SDS, and 0.6% tris base) supplemented with protease and phosphatase inhibitors (Sigma-Aldrich). Proteins were resolved by SDS-PAGE and transferred to PVDF membranes, as previously described (34, 42). Gonadal and inguinal adipose tissues were harvested from WT and adi14-3-3ζKO mice, fixed in 4% paraformaldehyde (Sigma-Aldrich), embedded in paraffin, and sectioned to 6 μm thickness.

Immunohistochemistry for perilipin (Cell Signaling Technology, Danvers, MA) or perilipin (Fitzgerald, Acton, MA) and 14-3-3ζ (Abcam, Toronto, ON) was performed with established protocols, and Cell Profiler (3.0.0) was used to determine adipocyte size, as previously described (34, 38). To image lipid droplets, mature 3T3-L1 adipocytes were seeded in chamber slides (ThermoFisher Scientific) coated with Poly-D-lysine (Sigma-Aldrich) after electroporation with either a scrambled, control (siCon) or siRNA against 14-3-3ζ, and cells were incubated with 0.5 μm Hoechst 33342 and 2 μm Bodipy 493/503 (ThermoFisher Scientific). Following successive washes with PBS, cells were fixed with 4% paraformaldehyde and cover-slipped after addition of ProLong Gold mounting medium (ThermoFisher Scientific). All images were taken with an Evos FL fluoresecent microscope (ThermoFisher Scientific). All antibodies and their concentrations are listed in Supplemental Table 2.

### Untargeted lipidomic analysis (LC-MS)

Methanol fixed, siRNA-transfected, differentiated 3T3-L1 cells were processed for lipid extraction and untargeted LC-MS lipidomics, as previously described (20). In brief, samples (0.50 μl to 1.82 μl), spiked with six internal standards: LPC 13:0, PC19:0/19:0, PC14:0/14:0, PS12:0/12:0, PG15:0/15:0 and PE17:0/17:0 (Avanti Polar Lipids Inc, Alabaster, USA), were analyzed using a 1290 Infinity HPLC coupled to a 6530 Accurate Mass Quadrupole-Time-of-Flight (QTOF) equipped with a dual ion spray electrosource (Agilent, Santa Clara, CA) and operated in the positive mode. Lipids were eluted on a Zorbax Eclipse plus column (C18, 2.1 x 100 mm, 1.8 μm, Agilent Technologies Inc.) over 83 minutes at 40 °C using a gradient of solvent A (0.2% formic acid and 10 mM ammonium formate in water) and B (0.2% formic acid and 5 mM ammonium formate in methanol/acetonitrile/methyl tert-butyl ether [MTBE], 55:35:10 [v/v/v]). MS data processing was achieved using Mass Hunter Qualitative Analysis software package (version B.06) and a bioinformatic script that we developed and encoded in both Perl and R languages to enable optimal MS data alignment between runs. This yielded a data set listing features with their mass (*m/z*), corrected signal intensity, and retention time. Lipid identification was achieved by alignment using an in-house reference database in which 498 lipids have previously been identified by MS/MS.

### Statistical analyses

Data are expressed as mean ± standard error (SE) and were analyzed by one- or two-way ANOVA or Student’s t-test. A p-value less than 0.05 was considered statistically significant. For lipidomic analysis, the output text file containing the processed data was imported into Mass Professional Pro software (version 12.6.1, Agilent Technologies Inc.) and independent testing of each feature was achieved using an unpaired student’s *t*-test followed by Benjamini-Hochberg correction. For selecting features discriminating siCon- vs. si14-3-3ζ-transfected cells, we have applied a corrected *P*-value as an estimation of False Discovery Rate (FDR) of < 5% and and a fold change (FC) >2 and <0.5.

## Results

### Adipocyte-specific deletion of 14-3-3ζ impairs lipolysis in mice

To investigate the *in vivo* contributions of 14-3-3ζ to lipolysis, we generated tamoxifen (TMX)-inducible, adipocyte-specific 14-3-3ζ knockout (adi14-3-3ζKO) mice whereby exon 4 of *Ywhaz* was deleted (Figure 1A). Quantitative PCR confirmed that *Ywhaz* expression was significantly decreased in gonadal white (gWAT) and inguinal white (iWAT) adipose tissues, and no effects on expression of the remaining 6 isoforms were detected (Figure 1B, Figure S1A). Following TMX treatment, no effects on body weights between adi14-3-3ζKO and Cre+ wild-type (WT) littermate controls were observed in male and female mice (Figure 1C, Figure S1B), and adipocyte-specific deletion of 14-3-3ζ did not affect glucose metabolism or insulin sensitivity, as determined by intraperitoneal glucose tolerance (IPGTT) and intraperitoneal insulin tolerance (ITT) tests (Figures 1D,E, Figure S1C,D). Additionally, no differences in circulating insulin were detected in male mice following 14-3-3ζ deletion in adipocytes (Figure 1F).

Given the presence of 14-3-3 protein binding sites in ATGL and HSL (1, 36), we sought to examine whether deletion of 14-3-3ζ in adipocytes would impact lipolysis. WT and adi14-3-3ζKO mice were fasted overnight and challenged with the β3-adrenergic agonist, CL-316,243. In male adi14-3-3ζKO mice, a significant reduction in plasma glycerol levels following intraperitoneal administration of CL-316,243 was detected (Figure 1G); however, no significant differences were observed in female adi14-3-3ζKO mice (Figure S1E). Our observation of impaired glycerol release in male adi14-3-3ζKO mice prompted us to further examine if the effect of 14-3-3ζ was specific to adipose tissue. Explant studies with gWAT were performed whereby gWAT from male WT and adi14-3-3ζKO mice were stimulated with isoproterenol, and significantly impaired isoproterenol-mediated FFA and glycerol release were detected from adi14-3-3ζKO gonadal white adipose tissue (gWAT) explants (Figure 1H). To examine the consequence of 14-3-3ζ over-expression on lipolysis, transgenic mice over-expressing a TAP-tagged human 14-3-3ζ molecule and WT littermate controls were injected with isoproterenol, and no differences were observed (Figure S2A,B). Collectively, these data demonstrate that only adipocyte-specific deletion of 14-3-3ζ is sufficient to impair lipolysis.

We previously reported that systemic deletion of 14-3-3ζ in mice caused significant reductions in the size and mass gWAT (34). However, it was not clear if these effects were cell-autonomous and specific to 14-3-3ζ deletion in adipocytes. In the present study, no differences in average size or distribution of adipocytes in iWAT from adi14-3-3ζKO mice were observed (Figure 2A-C). In contrast, adipocytes within gWAT were significantly smaller, and decreases and increases in the relative proportion of smaller (<200 μm) and larger (>500 μm) adipocytes, respectively, were observed (Figure 2D-F). When taken together, these findings suggest that the predominant effect of 14-3-3ζ deletion may be restricted to adipocytes within visceral depots like gonadal adipose tissue.

**Figure 2.**
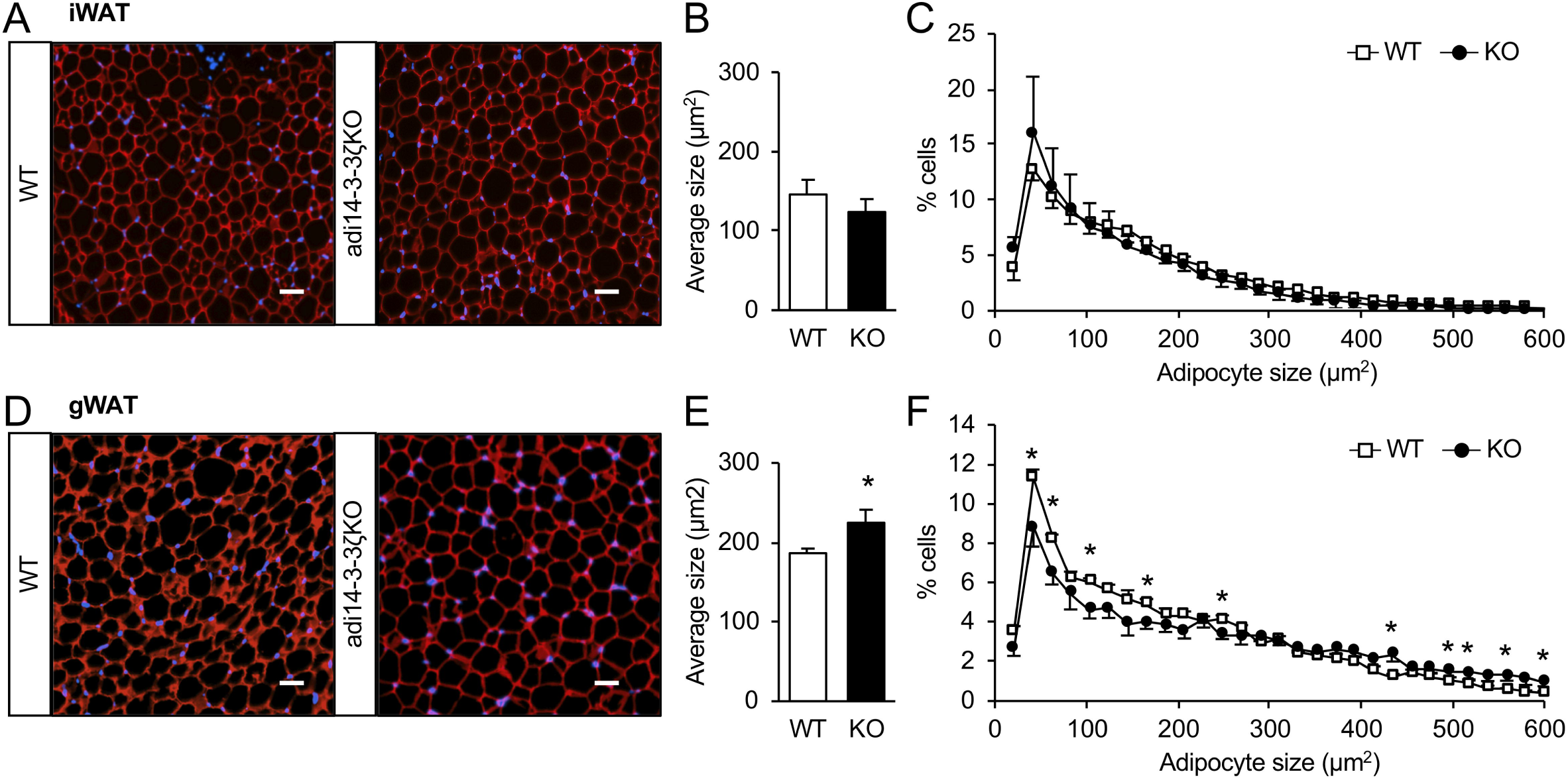
Loss of 14-3-3ζ in gonadal white adipocytes affects adipocyte size. **(A-F)** Immunofluorescent staining was used to examine adipocyte morphology (A, D), average size (B,E), and distrubtion (C,F) in inguinal white adipose tissue (iWAT, A-C) or gonadal white adipose tissue (gWAT, D-F) of male WT and KO mice. (Representative images of n=4 mice per group; scale bar= 200 μm; Student’s t-test to compare WT vs KO; *: p<0.05 when compared to WT). All values represent mean ± SE.

### 14-3-3ζ depletion impairs lipolysis in 3T3-L1 cells

To further understand how 14-3-3ζ could influence lipolysis, differentiated 3T3-L1 adipocytes were transfected with siRNA against 14-3-3ζ (Figure 3A,B). No off-target effects on the expression of the remaining 14-3-3 isoforms were detected (Figure 3C). Glycerol and FFA levels were measured in the supernatant of 14-3-3ζ-depleted 3T3-L1 adipocytes after the addition of lipolytic stimuli, and when compared to control cells (siCon), knockdown of 14-3-3ζ (si14-3-3ζ) significantly impaired lipolysis in response to isoproterenol, forskolin, and dibutyryl cAMP (dbcAMP) (Figure 3D). As each compound targets effectors at different points in the β3-adrenergic receptor pathway, these findings suggested that depletion of 14-3-3ζ affected a distal event that facilitates lipolysis. In contrast to what was observed with over-expression of 14-3-3ζ *in vivo*, transient transfection of mature 3T3-L1 adipocytes with plasmid to over-express 14-3-3ζ was associated with a potentiation in isoproterenol-mediated lipolysis (Figure S2C,D).

**Figure 3.**
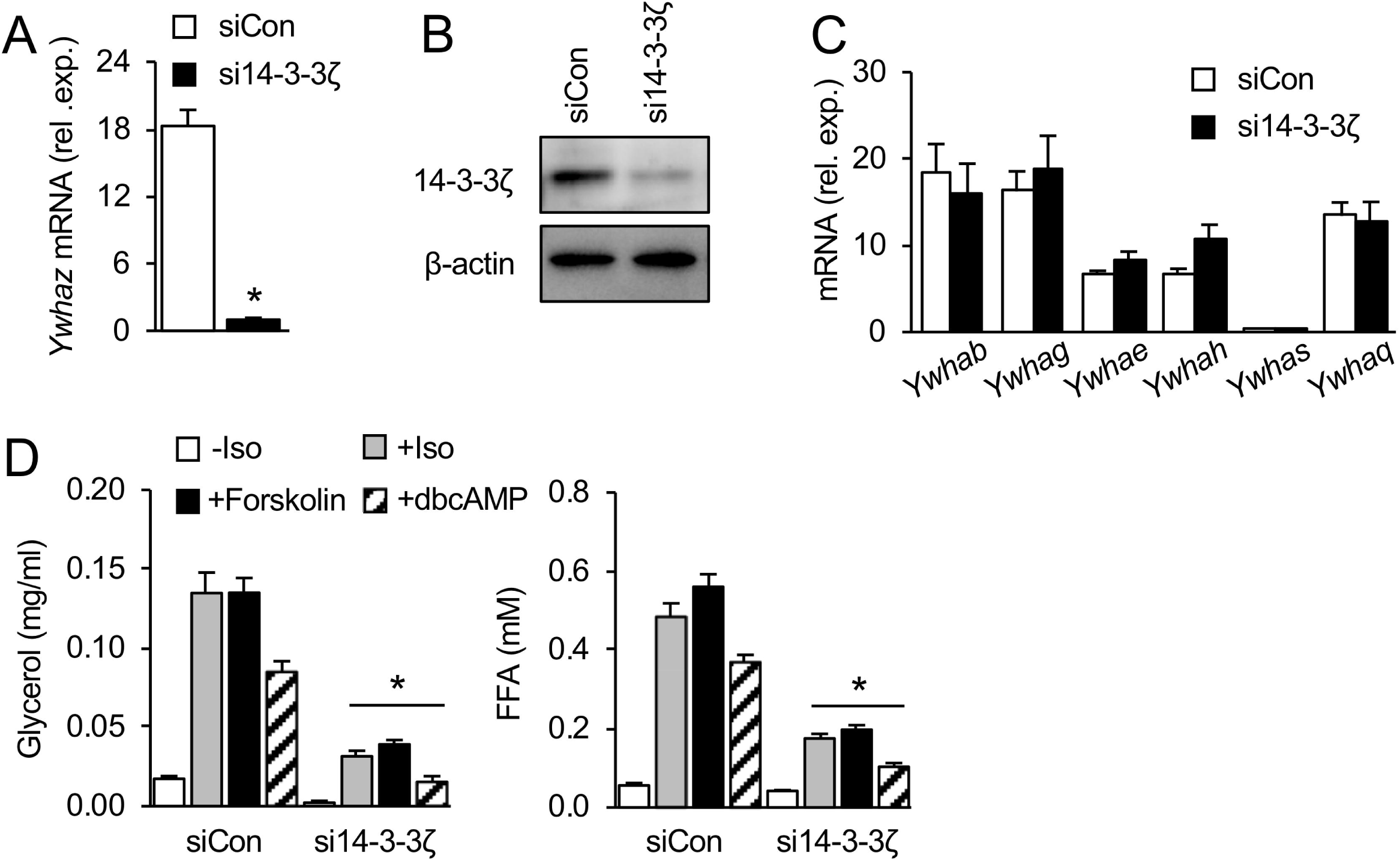
Depletion of 14-3-3ζ in mature 3T3-L1 adipocytes impairs lipolysis in response to mulitple stimuli. **(A**,**B)** Depletion of 14-3-3ζ with siRNA in mature 3T3-L1 adipocytes was confirmed by qPCR (A) (n=6 per group; Student’s t-test *= p<0.05 vs. siCon) and immunoblotting of cell lysates for 14-3-3ζ (B) (n=3 per condition). **(C)** Levels of *Ywhab, Ywhag, Ywhae, Ywhah, Ywhas* and *Ywhaq* (encoding 14-3-3β, 14-3-3γ, 14-3-3ε, 14-3-3η, 14-3-3σ, and 14-3-3θ, respectively) mRNA were measured in 14-3-3ζ-depleted 3T3-L1 adipocytes (n=6 per group). **(D)** Levels of glycerol (C) and FFA (D) released by siRNA-transfected 3T3-L1 adipocytes treated with isoproterenol (Iso, 1 μM), forskolin (10 μM) with IMBX (0.5 mM), or dibutyryl cAMP (dbcAMP, 1 mM) for 2 hours (n=6 per condition; two-way ANOVA, followed by Bonferroni t-tests; *= p<0.05 when compared to respective treatment in siCon-transfected cells). All values represent mean ± SE.

As 14-3-3ζ depletion impaired lipolysis, it prompted further examination of whether the activity of the β3-adrenergic signaling pathway was affected. We first looked at mRNA expression of the β-adrenergic receptor isoforms (*Adrb1, Adrb2*, and *Adrb3*) in mature, 14-3-3ζ-depleted 3T3-L1 adipocytes. While there were no changes in mRNA levels for *Adrb1* and *Adrb2*, a significant reduction in *Adrb3* mRNA levels, which is mainly expressed in adipocytes (49), was observed (Figure 4A). The attenuation of forskolin-mediated lipolysis suggested that defects in adenylyl cyclase activity could be occurring following 14-3-3ζ depletion (Figure 3D). However, no differences in the abilities of isoproterenol or forskolin to stimulate cAMP production were observed (Figure 4C), nor were significant differences in the expression of various phosphodiesterases, which degrade cAMP, detected between groups (Figure 4B). No differences in the activity of PKA, as assessed by HSL or CREB phosphorylation, was observed following 14-3-3ζ knockdown (Figure 4F); instead, significant reductions in mRNA and protein levels of *Atgl/*ATGL and *Hsl/*HSL were detected in 14-3-3ζ-depleted 3T3-L1 cells (Figure 4D,E), in addition to protein abundance of the PKA target, CREB (Figure 4E). Collectively, these findings demonstrate that the defect in lipolysis following 14-3-3ζ depletion cannot be solely attributed to impairments in the β3-adrenergic receptor-PKA signaling pathway.

**Figure 4.**
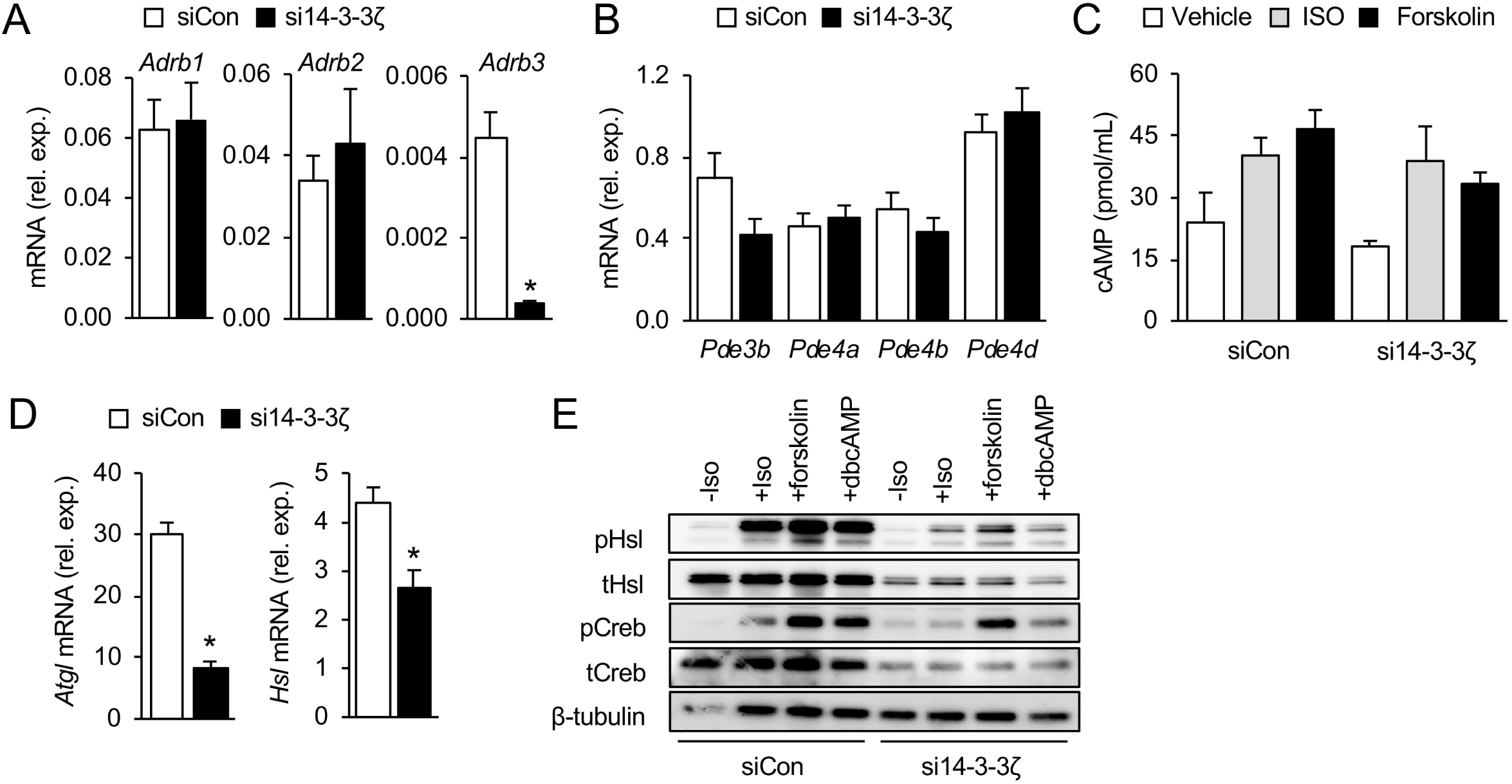
14-3-3ζ is required for the expression of lipolytic effectors. **(A**,**B)** Quantitative PCR was used to measure mRNA levels of β-adrenergic receptor (A) and phosphodiesterase (B) isoforms in 14-3-3ζ-depleted 3T3-L1 adipocytes (n=6 per group; Student’s t-test; *= p<0.05 when compared to siCon). **(C)** Intracellular cAMP levels of siRNA-transfected 3T3-L1 lysates treated with isoproterenol (ISO, 1 μM) or forskolin (20 μM) for 1 hour (n=6-7 per group). **(D)** mRNA levels of *Atgl* and *Hsl* in mature 3T3-L1 cells following siRNA-mediated knockdown of 14-3-3ζ (n=6 per group; Student’s t-test; *= p<0.05 when compared to siCon). **(E)** Immunoblotting for phosphorylated and total forms of HSL and CREB in siRNA-transfected 3T3-L1 lysates treated with isoproterenol (ISO, 1 μM), forskolin (10 μM) with IMBX (0.5 mM), or dbCAMP (1 mM) (n=6 per group). All values represent mean ± SE.

### Deletion of 14-3-3ζ in primary white adipocytes reduces the expression of mature adipocyte markers

To examine whether 14-3-3ζ deletion in primary adipocytes could cause similar reductions in the expression of lipases, quantitative PCR and immunoblotting was performed. Unlike what was observed in 3T3-L1 cells, mRNA levels of *Atgl* and *Hsl* were not affected by deletion of 14-3-3ζ in primary adipocytes (Figure 5A,B), but immunoblotting for HSL revealed marked reductions in protein abundance in gWAT from adi14-3-3ζKO mice (Figure 5D). Moreover, PPARγ2 protein abundance was reduced in gWAT from adi14-3-3ζKO mice, which suggests a loss of adipocyte identity or maturity (Figure 5D). No differences in total TAGs were observed in gWAT or iWAT from WT and adi14-3-3ζKO mice (Figure 5E), but genes associated with *de novo* triglyceride synthesis were significantly reduced only in gWAT of adi14-3-3-ζKO mice (Figure 5F,G). Furthermore, examination of genes associated with adipocyte maturity was performed, and significant reductions in mature adipocyte genes, such as *Adipoq, Fabp4*, and *Retn*, could be detected in both iWAT and gWAT from adi14-3-3ζKO animals (Figure 5H,I). Despite the decreased levels of *Adipoq* mRNA in iWAT and gWAT, no differences in circulating adiponectin levels were observed, in addition to plasma leptin levels (Figure 5J,K).

**Figure 5.**
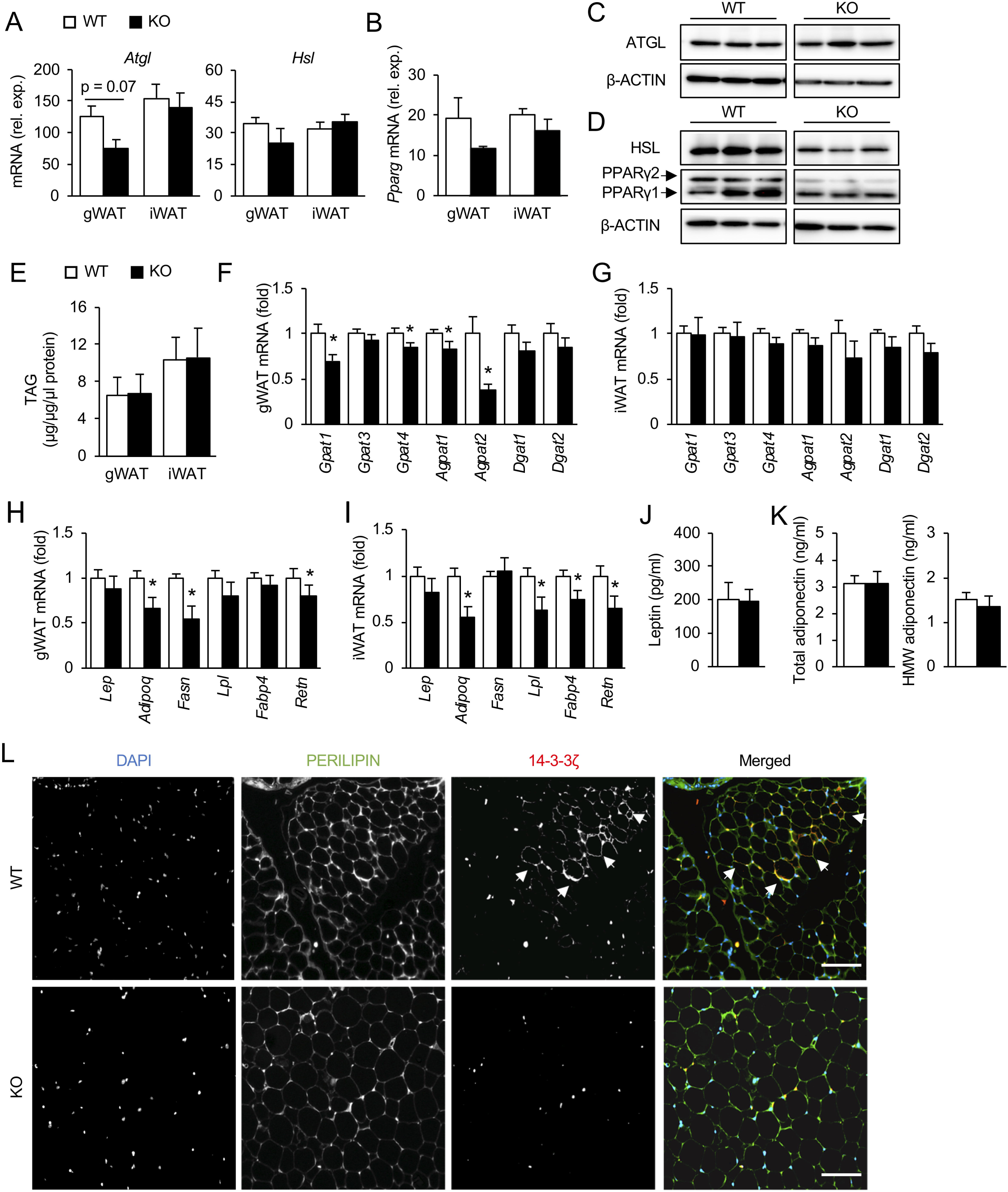
Adipocyte-specific deletion of 14-3-3ζ decreases expression of genes associated with adipocyte maturity *in vivo*. **(A**,**B)** Quantitative PCR was used to measure mRNA levels of *Atgl* and *Hsl* (A) and *Pparg* (B) in gonadal (gWAT) and inguinal (iWAT) adipose tissues of male adi14-3-3ζKO mice (n=6 per genotype). **(C-D)** Immunoblotting was used to measure total levels of ATGL (C) and HSL and PPARγ (D) in gWAT of male adi14-3-3ζKO mice (representative of n=6 per genotype). **(E)** Total triacylglycerols (TAGs) were measured in gWAT and iWAT of WT and adi14-3-3ζKO (KO) mice. Data were normalized to total protein of each sample (n=6 per genotype). **(F-I)** Genes associated with *de novo* triglyceride synthesis (F,G) and adipocyte maturity (H,I) were measured in gWAT (F,H) and iWAT (G,I) of WT or adi14-3-3ζKO mice (n=6 per genotype; Student’s t-test; *: p<0.05 when compared to WT mice). **(J**,**K)** Leptin (J) and adiponectin (K) levels were determined from WT and adi14-3-3ζKO mouse plasma collected after an overnight fast (n=6 per genotype). **(L)** Immunohistochemistry was performed on paraffin-embedded gWAT sections of WT and adi14-3-3ζKO mice. Arrowheads denote adipocytes that display 14-3-3ζ immunoreactivity. (representative images of n=4 mice per group). All values represent mean ± SE.

The observed small, but significant impairments in *ex vivo* lipolysis (Figure 1H), changes in the size and distribution of gonadal white adipocytes (Figure 2D,F), and reductions in adipocyte maturity genes (Figure 5F,H), suggest the possibility that not all adipocytes within gWAT are affected by 14-3-3ζ deletion. Indeed, using immunohistochemistry, we confirmed the existence of a subset of adipocytes within gWAT that express 14-3-3ζ. These 14-3-3ζ-expressing adipocytes were undetectable in adi14-3-3ζKO mice (Figure 5L). Thus, the phenotype observed in adi14-3-3ζKO mice with respect to lipolysis may be attributed to a specific subset of adipocytes within gWAT that have lost their maturity.

### Depletion of 14-3-3ζ promotes adipocyte immaturity *in vitro*

To further characterize how reductions in 14-3-3ζ could affect adipocyte maturity, 14-3-3ζ-depleted mature 3T3-L1 adipocytes were used. Initially, Oil Red-O incorporation was used as a surrogate measure of lipid content, and knockdown of 14-3-3ζ was found to significantly reduce lipids present in 3T3-L1 cells and was associated with marked decreases in cells containing lipid droplets (Figure 6A,B). This was also confirmed by Bodipy 493/503 imaging and biochemical measurements of total TAGs, both of which demonstrated significant decreases in the number of Bodipy-labelled lipid droplets and TAG content, respectively (Figure 6C,D). The impact of 14-3-3ζ depletion on TAG content was more pronounced in 3T3-L1 cells than in our *in vivo* model (Figure 6D vs Figure 5E), but this could be due to the heterogeneous expression of 14-3-3ζ within gWAT (Figure 5L). Similar to what was observed in adi14-3-3ζKO adipocytes, genes associated with *de novo* triglyceride synthesis and adipocyte maturity were significantly decreased (Figure 6E,F).

**Figure 6.**
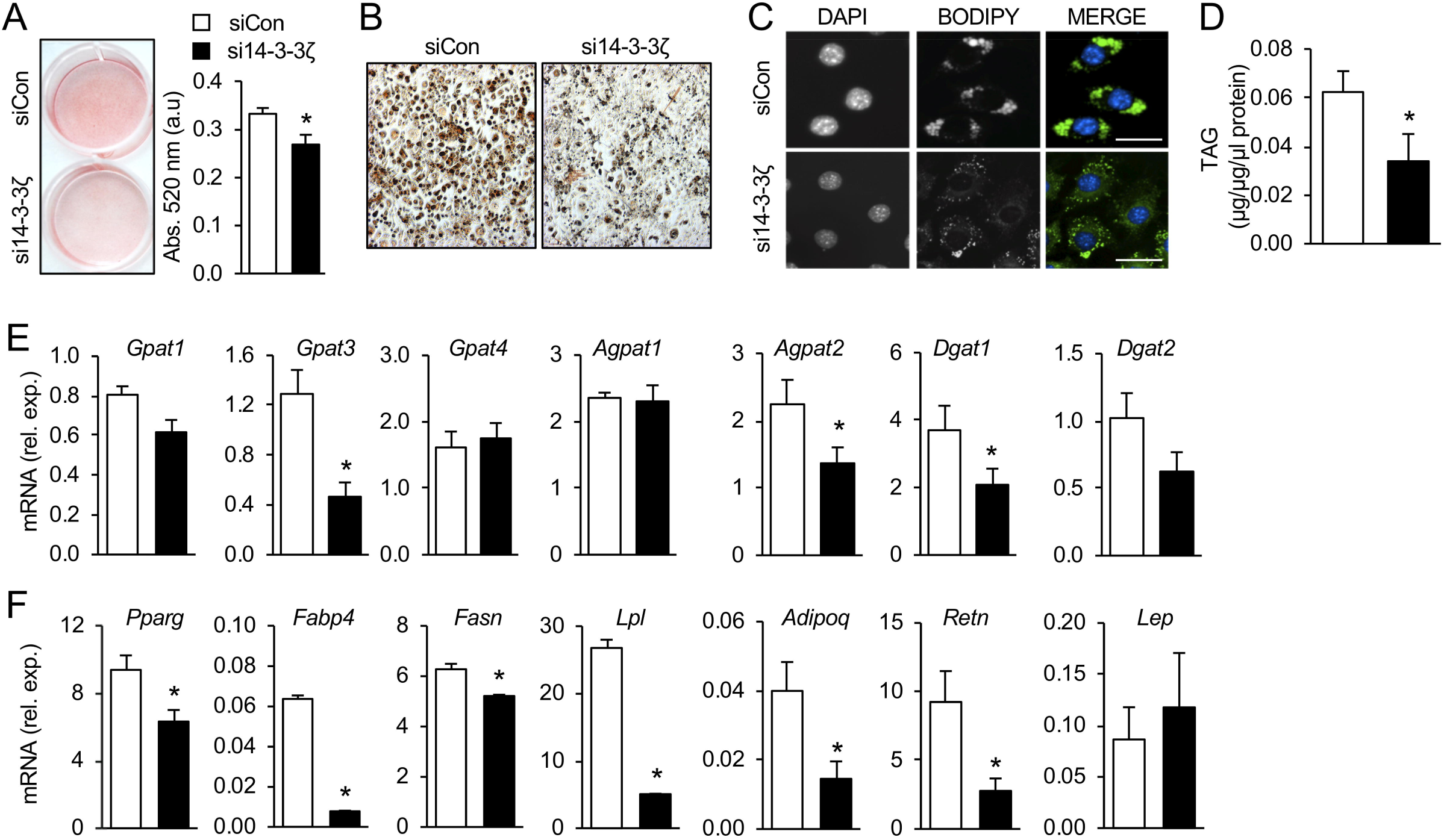
Knockdown of 14-3-3ζ by siRNA promotes dedifferentiation *in vitro*. **(A)** Oil Red-O incorporation was used to assess triglyceride levels in siCon- and si14-3-3ζ-transfected mature 3T3-L1 adipocytes (n=4; Student’s t-test; *= p<0.05 siCon vs si14-3-3ζ). **(B)** Visualization of Oil Red-O incorporated control (siCon) and 14-3-3ζ-depleted (si14-3-3ζ) 3T3-L1 adipocytes by light microscopy (representative images of n=4 per treatment). **(C)** Lipid droplets were visualized by immunofluorescent imaging of incorporated Bodipy 493/503. Hoechst 33342 was used to visualize nuclei (representative image of transfected cells, scale bar= 25 μm). **(D)** Total triglycerides were measured in mature 3T3-L1 adipocytes transfected with a scrambled control (siCon) or siRNA against 14-3-3ζ (si14-3-3ζ) (n=6; Student’s t-test; *= p<0.05 siCon vs si14-3-3ζ). **(E**,**F)** Following transfection with siRNA against 14-3-3ζ (si14-3-3ζ) or a scrambled control (siCon), mRNA was isolated, followed by quantitative PCR for genes associated with *de novo* triglyceride synthesis (E) or adipocyte maturity (F) (n=6 per group, Student’s t-test; *: p<0.05 when compared to siCon). All values represent mean ± SE.

### Knockdown of 14-3-3ζ is associated with distinct changes in the 3T3-L1 lipidome

During 3T3-L1 adipocyte differentiation, significant changes in the lipidome, including specific TAG species and sphingomyelins, have been observed (33). Conversely, de-differentiation of primary, mature adipocytes is associated with changes in the abundance of specific lipid species (4). When taken together, these findings suggest that identification of specific lipidomic signatures could be an alternative approach to assess adipocyte maturity. To this end, we used untargeted LC-MS lipidomics to interrogate lipid species that are changed upon depletion of 14-3-3ζ in mature 3T3-L1 cells and to confirm if reductions in 14-3-3ζ can generate a lipidomic signature associated with adipocyte dedifferentiation (Figure 7A). A total of 2535 species were detected, of which 549 passed the stringent selection criteria of FDR<5% and FC >2 or < 0.5. A total of 139 annotated lipids from different lipid (sub)classes were significantly changed following 14-3-3ζ depletion (Figure 7B). For example, knockdown of 14-3-3ζ predominantly enriched monoacylglycerophosphocholines (LPC) and diacylglycerophosphocholines (PC), whereas diacylglycerophosphoethanolamines (PE and PE(O)) were reduced (Figure 7B, Table 1). The vast majority of altered lipid species (73/139) were triglycerides, and 14-3-3ζ depletion primarily decreased triglycerides species with lower acyl carbon chains and lower degrees of carbon bond saturation (Figure 7C).

**Table 1.** List of lipid species that were significantly up- and down-regulated following 14-3-3ζ depletion.

**Down regulated entities**
| Compound | ionisation<br>mode | Masse | m/z | RT | Adduct | p | p (Corr) | FC (abs) | Regulation<br>(SiZeta vs SiCon) | FC |
| --- | --- | --- | --- | --- | --- | --- | --- | --- | --- | --- |
| LPC24:0 | POS | 607.4558 | 608.4636 | 16.71 | [M+H]<sup>+</sup> | 1.22E-03 | 5.22E-03 | 2.66 | down | -2.66 |
| PE(14:0_22:6) | POS | 735.4838 | 736.4916 | 17.99 | [M+H]<sup>+</sup> | 5.86E-04 | 3.08E-03 | 2.12 | down | -2.12 |
| PE(16:0_18:1) | POS | 717.5301 | 718.5379 | 26.12 | [M+H]<sup>+</sup> | 8.47E-06 | 1.83E-04 | 2.05 | down | -2.05 |
| PE(34:2) | POS | 715.5146 | 716.5224 | 22.86 | [M+H]<sup>+</sup> | 4.66E-06 | 1.17E-04 | 2.95 | down | -2.95 |
| PE(36:2) | POS | 743.5465 | 744.5543 | 26.52 | [M+H]<sup>+</sup> | 1.28E-05 | 2.38E-04 | 2.27 | down | -2.27 |
| PE(O-18:0/18:1) | POS | 731.5832 | 732.591 | 32.48 | [M+H]<sup>+</sup> | 1.67E-04 | 1.33E-03 | 3.12 | down | -3.12 |
| PE(O-18:1/18:1) | POS | 729.5661 | 730.5739 | 31.82 | [M+H]<sup>+</sup> | 1.38E-03 | 5.72E-03 | 3.71 | down | -3.71 |
| PE(O-18:2/18:2) | POS | 725.5349 | 726.5427 | 25.52 | [M+H]<sup>+</sup> | 5.78E-05 | 6.54E-04 | 8.72 | down | -8.72 |
| PE(O-36:3) | POS | 727.551 | 728.5588 | 28.82 | [M+H]<sup>+</sup> | 1.03E-04 | 9.63E-04 | 3.35 | down | -3.35 |
| PE(O-34:3) | POS | 699.5192 | 700.527 | 24.95 | [M+H]<sup>+</sup> | 4.85E-05 | 5.90E-04 | 3.89 | down | -3.89 |
| PE(O-34:2) | POS | 701.5323 | 702.5401 | 27.78 | [M+H]<sup>+</sup> | 1.13E-04 | 1.02E-03 | 3.99 | down | -3.99 |
| PI(16:0_20:4) | POS | 875.5525 | 876.5603 | 18.17 | [M+NH4]<sup>+</sup> | 9.78E-06 | 2.01E-04 | 2.21 | down | -2.21 |
| TG(16:0_15:0_18:1) | POS | 840.722 | 841.7298 | 55.85 | [M+Na]<sup>+</sup> | 1.45E-04 | 1.22E-03 | 2.14 | down | -2.14 |
| TG(16:0_16:0_18:1) | POS | 854.7393 | 855.7471 | 59.61 | [M+Na]<sup>+</sup> | 1.96E-04 | 1.47E-03 | 2.37 | down | -2.37 |
| TG(18:2_18:2_12:0) | POS | 815.7022 | 816.71 | 42.55 | [M+NH4]<sup>+</sup> | 5.45E-05 | 6.25E-04 | 3.37 | down | -3.37 |
| TG(18:3_18:1_12:0) | POS | 815.7027 | 816.7105 | 42.79 | [M+NH4]<sup>+</sup> | 1.35E-04 | 1.15E-03 | 2.13 | down | -2.13 |
| TG 39:0 | POS | 697.6248 | 698.6326 | 40.4 | [M+NH4]<sup>+</sup> | 1.80E-06 | 6.32E-05 | 2.98 | down | -2.98 |
| TG 39:0 | POS | 697.6248 | 698.6326 | 40 | [M+NH4]<sup>+</sup> | 1.91E-04 | 1.44E-03 | 2.11 | down | -2.11 |
| TG 39:1 | POS | 695.6087 | 696.6165 | 38.62 | [M+NH4]<sup>+</sup> | 4.76E-05 | 5.88E-04 | 4.14 | down | -4.14 |
| TG 40:1 | POS | 709.6248 | 710.6326 | 39.55 | [M+NH4]<sup>+</sup> | 7.19E-06 | 1.66E-04 | 3.26 | down | -3.26 |
| TG 40:2 | POS | 707.6085 | 708.6163 | 37.94 | [M+NH4]<sup>+</sup> | 3.30E-05 | 4.59E-04 | 3.54 | down | -3.54 |
| TG 41:0 | POS | 725.6559 | 726.6637 | 42.74 | [M+NH4]<sup>+</sup> | 1.11E-07 | 9.60E-06 | 2.64 | down | -2.64 |
| TG 41:1 | POS | 723.6399 | 724.6477 | 40.56 | [M+NH4]<sup>+</sup> | 4.65E-06 | 1.17E-04 | 3.58 | down | -3.58 |
| TG 41:2 | POS | 721.6242 | 722.632 | 38.8 | [M+NH4]<sup>+</sup> | 2.38E-05 | 3.60E-04 | 4.03 | down | -4.03 |
| TG 42:2 | POS | 735.6393 | 736.6471 | 39.72 | [M+NH4]<sup>+</sup> | 1.24E-05 | 2.38E-04 | 3.69 | down | -3.69 |
| TG 43:0 | POS | 753.6882 | 754.696 | 45.99 | [M+NH4]<sup>+</sup> | 1.22E-05 | 2.38E-04 | 2.17 | down | -2.17 |
| TG 43:1 | POS | 751.671 | 752.6788 | 42.51 | [M+NH4]<sup>+</sup> | 6.52E-04 | 3.29E-03 | 3.08 | down | -3.08 |
| TG 43:1 | POS | 751.6709 | 752.6787 | 42.95 | [M+NH4]<sup>+</sup> | 2.17E-05 | 3.48E-04 | 2.87 | down | -2.87 |
| TG 43:2 | POS | 749.6569 | 750.6647 | 41.14 | [M+NH4]<sup>+</sup> | 3.18E-03 | 1.06E-02 | 3.29 | down | -3.29 |
| TG 43:2 | POS | 749.6555 | 750.6633 | 40.7 | [M+NH4]<sup>+</sup> | 7.98E-06 | 1.80E-04 | 3.21 | down | -3.21 |
| TG 43:3 | POS | 747.6403 | 748.6481 | 39.08 | [M+NH4]<sup>+</sup> | 2.18E-05 | 3.48E-04 | 3.55 | down | -3.55 |
| TG 44:1 | POS | 765.6875 | 766.6953 | 45.28 | [M+NH4]<sup>+</sup> | 2.44E-04 | 1.69E-03 | 17.67 | down | -17.67 |
| TG 44:1 | POS | 765.6871 | 766.6949 | 43.96 | [M+NH4]<sup>+</sup> | 9.51E-04 | 4.28E-03 | 7.54 | down | -7.54 |
| TG 44:2 | POS | 763.6717 | 764.6795 | 42.39 | [M+NH4]<sup>+</sup> | 7.08E-04 | 3.48E-03 | 3.14 | down | -3.14 |
| TG 44:2 | POS | 763.6706 | 764.6784 | 41.86 | [M+NH4]<sup>+</sup> | 6.05E-05 | 6.62E-04 | 2.55 | down | -2.55 |
| TG 44:3 | POS | 761.6561 | 762.6639 | 40.25 | [M+NH4]<sup>+</sup> | 5.36E-05 | 6.25E-04 | 3.25 | down | -3.25 |
| TG 44:3 | POS | 761.6557 | 762.6635 | 39.94 | [M+NH4]<sup>+</sup> | 1.57E-04 | 1.29E-03 | 3.07 | down | -3.07 |
| TG 45:1 | POS | 779.7032 | 780.711 | 45.64 | [M+NH4]<sup>+</sup> | 1.26E-04 | 1.09E-03 | 4.46 | down | -4.46 |
| TG 45:2 | POS | 777.6873 | 778.6951 | 43.79 | [M+NH4]<sup>+</sup> | 1.22E-03 | 5.22E-03 | 2.64 | down | -2.64 |
| TG 45:3 | POS | 775.6706 | 776.6784 | 40.93 | [M+NH4]<sup>+</sup> | 4.04E-04 | 2.44E-03 | 2.47 | down | -2.47 |
| TG 45:4 | POS | 773.6569 | 774.6647 | 39.49 | [M+NH4]<sup>+</sup> | 1.04E-02 | 2.71E-02 | 2.97 | down | -2.97 |
| TG 46:1 | POS | 793.7186 | 794.7264 | 47.63 | [M+NH4]<sup>+</sup> | 4.13E-04 | 2.46E-03 | 2.09 | down | -2.09 |
| TG 46:3 | POS | 789.6866 | 790.6944 | 42.56 | [M+NH4]<sup>+</sup> | 2.42E-03 | 8.54E-03 | 2.35 | down | -2.35 |
| TG 46:4 | POS | 787.6688 | 788.6766 | 40.44 | [M+NH4]<sup>+</sup> | 6.79E-05 | 7.14E-04 | 3.65 | down | -3.65 |
| TG 46:4 | POS | 787.6686 | 788.6764 | 40.98 | [M+NH4]<sup>+</sup> | 3.68E-03 | 1.19E-02 | 2.77 | down | -2.77 |
| TG 47:1 | POS | 807.7336 | 808.7414 | 51.35 | [M+NH4]<sup>+</sup> | 7.30E-04 | 3.56E-03 | 2.01 | down | -2.01 |
| TG 47:2 | POS | 805.7191 | 806.7269 | 45.9 | [M+NH4]<sup>+</sup> | 2.11E-03 | 7.74E-03 | 2.70 | down | -2.70 |
| TG 47:3 | POS | 803.7031 | 804.7109 | 44.01 | [M+NH4]<sup>+</sup> | 1.54E-03 | 6.18E-03 | 2.41 | down | -2.41 |
| TG 47:4 | POS | 801.6871 | 802.6949 | 41.54 | [M+NH4]<sup>+</sup> | 7.89E-05 | 7.99E-04 | 3.21 | down | -3.21 |
| TG 48:3 | POS | 817.7181 | 818.7259 | 45.7 | [M+NH4]<sup>+</sup> | 6.81E-03 | 1.95E-02 | 2.68 | down | -2.68 |
| TG 49:3 | POS | 831.7349 | 832.7427 | 47.66 | [M+NH4]<sup>+</sup> | 1.72E-03 | 6.64E-03 | 3.87 | down | -3.87 |
| TG 49:3 | POS | 859.7577 | 860.7655 | 49.06 | [M+NH4]<sup>+</sup> | 1.02E-02 | 2.67E-02 | 2.11 | down | -2.11 |
| TG 49:4 | POS | 829.7197 | 830.7275 | 44.25 | [M+NH4]<sup>+</sup> | 1.33E-05 | 2.47E-04 | 2.99 | down | -2.99 |
| TG 49:4 | POS | 829.7191 | 830.7269 | 45.08 | [M+NH4]<sup>+</sup> | 9.04E-04 | 4.16E-03 | 2.98 | down | -2.98 |
| TG 49:4 | POS | 829.719 | 830.7268 | 44.6 | [M+NH4]<sup>+</sup> | 1.10E-04 | 1.00E-03 | 2.81 | down | -2.81 |
| TG 49:4 | POS | 829.7197 | 830.7275 | 43.91 | [M+NH4]<sup>+</sup> | 6.31E-05 | 6.70E-04 | 2.75 | down | -2.75 |
| TG 49:5 | POS | 827.7039 | 828.7117 | 42 | [M+NH4]<sup>+</sup> | 7.08E-04 | 3.48E-03 | 4.50 | down | -4.50 |
| TG 50:4 | POS | 843.7345 | 844.7423 | 45.96 | [M+NH4]<sup>+</sup> | 2.17E-05 | 3.48E-04 | 2.38 | down | -2.38 |
| TG 50:4 | POS | 843.734 | 844.7418 | 46.92 | [M+NH4]<sup>+</sup> | 1.10E-03 | 4.86E-03 | 2.14 | down | -2.14 |
| TG 50:5 | POS | 841.7172 | 842.725 | 43.32 | [M+NH4]<sup>+</sup> | 2.49E-05 | 3.74E-04 | 4.04 | down | -4.04 |
| TG 51:5 | POS | 855.735 | 856.7428 | 44.86 | [M+NH4]<sup>+</sup> | 4.58E-04 | 2.64E-03 | 4.46 | down | -4.46 |
| TG 51:5 | POS | 855.7338 | 856.7416 | 45.32 | [M+NH4]<sup>+</sup> | 2.67E-04 | 1.82E-03 | 2.76 | down | -2.76 |
| TG 53:4 | POS | 885.777 | 886.7848 | 47.44 | [M+NH4]<sup>+</sup> | 8.33E-03 | 2.27E-02 | 2.46 | down | -2.46 |
| TG 53:5 | POS | 883.7652 | 884.773 | 48.15 | [M+NH4]<sup>+</sup> | 3.14E-04 | 2.05E-03 | 2.20 | down | -2.20 |

**UP regulated**
| Compound | ionisation r | Masse | m/z | RT | Adduct | p | p (Corr) | FC (abs) | Regulation (SiZeta | FC |
| --- | --- | --- | --- | --- | --- | --- | --- | --- | --- | --- |
| LPC14:0-a | POS | 467.2995 | 468.3073 | 6.7 | [M+H]<sup>+</sup> | 2.77E-03 | 9.54E-03 | 3.82 | Up | 3.82 |
| LPC15:0 | POS | 481.3152 | 482.323 | 7.92 | [M+H]<sup>+</sup> | 1.58E-02 | 3.82E-02 | 2.06 | Up | 2.06 |
| LPC18:1 bc | POS | 521.3474 | 522.3552 | 9.49 | [M+H]<sup>+</sup> | 8.11E-03 | 2.22E-02 | 2.25 | Up | 2.25 |
| LPC19:0-b | POS | 537.3784 | 538.3862 | 11.59 | [M+H]<sup>+</sup> | 2.68E-06 | 8.10E-05 | 2.56 | Up | 2.56 |
| LPC20:3-b | POS | 545.3484 | 546.3562 | 9.04 | [M+H]<sup>+</sup> | 9.24E-03 | 2.48E-02 | 3.28 | Up | 3.28 |
| LPC20:4 b | POS | 543.3313 | 544.3391 | 8.16 | [M+H]<sup>+</sup> | 5.10E-05 | 6.06E-04 | 13.05 | Up | 13.05 |
| LPC20:5 | POS | 541.3151 | 542.3229 | 6.98 | [M+H]<sup>+</sup> | 3.84E-04 | 2.36E-03 | 8.83 | Up | 8.83 |
| LPC22:5 | POS | 569.3492 | 570.357 | 8.69 | [M+H]<sup>+</sup> | 6.94E-04 | 3.44E-03 | 6.23 | Up | 6.23 |
| LPC22:6-a | POS | 567.3307 | 568.3385 | 7.74 | [M+H]<sup>+</sup> | 1.06E-02 | 2.76E-02 | 3.23 | Up | 3.23 |
| LPC22:6-b | POS | 567.331 | 568.3388 | 8.1 | [M+H]<sup>+</sup> | 2.02E-04 | 1.51E-03 | 6.64 | Up | 6.64 |
| LPE20:4-b | POS | 501.2852 | 502.293 | 8.37 | [M+H]<sup>+</sup> | 1.85E-03 | 7.01E-03 | 4.90 | Up | 4.90 |
| LPE22:6-b | POS | 525.2839 | 526.2917 | 8.3 | [M+H]<sup>+</sup> | 3.46E-03 | 1.13E-02 | 4.73 | Up | 4.73 |
| PC(15:0_20:4) | POS | 767.5452 | 768.553 | 20.38 | [M+H]<sup>+</sup> | 9.42E-09 | 1.98E-06 | 4.45 | Up | 4.45 |
| PC(15:0_22:6) | POS | 791.5451 | 792.5529 | 19.75 | [M+H]<sup>+</sup> | 4.96E-05 | 5.97E-04 | 2.32 | Up | 2.32 |
| PC(16:0_15:0) | POS | 719.546 | 720.5538 | 22.71 | [M+H]<sup>+</sup> | 2.04E-06 | 6.87E-05 | 2.65 | Up | 2.65 |
| PC(16:0_16:0) | POS | 733.5607 | 734.5685 | 24.62 | [M+H]<sup>+</sup> | 2.38E-05 | 3.60E-04 | 2.50 | Up | 2.50 |
| PC(16:0_20:4n-3) | POS | 781.5604 | 782.5682 | 22.13 | [M+H]<sup>+</sup> | 9.12E-09 | 1.98E-06 | 4.74 | Up | 4.74 |
| PC(16:0_20:5) | POS | 779.5448 | 780.5526 | 19.99 | [M+H]<sup>+</sup> | 6.70E-08 | 7.05E-06 | 3.15 | Up | 3.15 |
| PC(16:0_22:4) | POS | 809.5916 | 810.5994 | 24.9 | [M+H]<sup>+</sup> | 6.60E-05 | 6.98E-04 | 2.71 | Up | 2.71 |
| PC(16:0_22:5) | POS | 807.5757 | 808.5835 | 23.56 | [M+H]<sup>+</sup> | 3.26E-08 | 4.65E-06 | 3.75 | Up | 3.75 |
| PC(17:0_20:4n-3) | POS | 795.5773 | 796.5851 | 23.99 | [M+H]<sup>+</sup> | 3.34E-09 | 1.98E-06 | 4.15 | Up | 4.15 |
| PC(17:0_22:6) | POS | 819.5757 | 820.5835 | 23.26 | [M+H]<sup>+</sup> | 7.25E-07 | 3.37E-05 | 2.14 | Up | 2.14 |
| PC(17:1_20:4) | POS | 793.5606 | 794.5684 | 20.99 | [M+H]<sup>+</sup> | 1.85E-04 | 1.42E-03 | 3.03 | Up | 3.03 |
| PC(18:0_20:4) | POS | 809.5941 | 810.6019 | 25.9 | [M+H]<sup>+</sup> | 4.05E-08 | 5.16E-06 | 5.33 | Up | 5.33 |
| PC(18:0_22:5n-3) | POS | 835.6064 | 836.6142 | 26.29 | [M+H]<sup>+</sup> | 8.29E-09 | 1.98E-06 | 2.98 | Up | 2.98 |
| PC(18:0_22:6) | POS | 833.5905 | 834.5983 | 25.12 | [M+H]<sup>+</sup> | 3.68E-08 | 4.95E-06 | 2.72 | Up | 2.72 |
| PC(18:1_22:5) | POS | 833.5923 | 834.6001 | 23.08 | [M+H]<sup>+</sup> | 1.21E-07 | 9.80E-06 | 2.83 | Up | 2.83 |
| PC(19:0_20:4) | POS | 823.6086 | 824.6164 | 27.85 | [M+H]<sup>+</sup> | 3.77E-05 | 4.99E-04 | 2.04 | Up | 2.04 |
| PC(20:1_22:6) | POS | 859.6084 | 860.6162 | 25.52 | [M+H]<sup>+</sup> | 2.08E-04 | 1.53E-03 | 2.23 | Up | 2.23 |
| PC(22:3_18:0) | POS | 839.6394 | 840.6472 | 31.55 | [M+H]<sup>+</sup> | 5.87E-05 | 6.55E-04 | 2.91 | Up | 2.91 |
| PC(18:1_22:6) | POS | 831.5771 | 832.5849 | 21.97 | [M+H]<sup>+</sup> | 4.43E-07 | 2.66E-05 | 2.59 | Up | 2.59 |
| PC(36:5) | POS | 779.5452 | 780.553 | 19.35 | [M+H]<sup>+</sup> | 3.05E-07 | 2.00E-05 | 3.22 | Up | 3.22 |
| PC(36:6) | POS | 777.529 | 778.5368 | 17.48 | [M+H]<sup>+</sup> | 3.23E-07 | 2.06E-05 | 2.19 | Up | 2.19 |
| PC(37:5) | POS | 793.5608 | 794.5686 | 21.73 | [M+H]<sup>+</sup> | 1.90E-07 | 1.44E-05 | 2.90 | Up | 2.90 |
| PC(38:3) | POS | 811.6094 | 812.6172 | 27.51 | [M+H]<sup>+</sup> | 2.61E-07 | 1.76E-05 | 2.66 | Up | 2.66 |
| PC(38:4)-OH | POS | 825.5867 | 826.5945 | 19.27 | [M+H]<sup>+</sup> | 2.23E-03 | 8.06E-03 | 2.23 | Up | 2.23 |
| PC(38:4) | POS | 809.5927 | 810.6005 | 24.21 | [M+H]<sup>+</sup> | 8.77E-06 | 1.85E-04 | 2.63 | Up | 2.63 |
| PC(38:4) | POS | 807.5778 | 808.5856 | 22.62 | [M+H]<sup>+</sup> | 8.51E-10 | 1.98E-06 | 4.94 | Up | 4.94 |
| PC(38:6) | POS | 805.5602 | 806.568 | 20.5 | [M+H]<sup>+</sup> | 7.80E-08 | 7.71E-06 | 4.47 | Up | 4.47 |
| PC(39:5) | POS | 821.5928 | 822.6006 | 24.43 | [M+H]<sup>+</sup> | 9.83E-08 | 9.15E-06 | 3.41 | Up | 3.41 |
| PC(39:6) | POS | 819.5757 | 820.5835 | 21.41 | [M+H]<sup>+</sup> | 4.42E-05 | 5.60E-04 | 2.06 | Up | 2.06 |
| PC(39:6) | POS | 819.5765 | 820.5843 | 22.18 | [M+H]<sup>+</sup> | 1.86E-04 | 1.42E-03 | 2.88 | Up | 2.88 |
| PC(40:4) | POS | 837.6233 | 838.6311 | 28.73 | [M+H]<sup>+</sup> | 1.02E-05 | 2.04E-04 | 2.41 | Up | 2.41 |
| PC(40:6) | POS | 833.5928 | 834.6006 | 23.93 | [M+H]<sup>+</sup> | 2.25E-03 | 8.09E-03 | 3.65 | Up | 3.65 |
| PC(O-16:0/20:4) | POS | 767.5803 | 768.5881 | 24.2 | [M+H]<sup>+</sup> | 1.08E-07 | 9.60E-06 | 4.43 | Up | 4.43 |
| PC(O-16:1/22:5) | POS | 791.5811 | 792.5889 | 24 | [M+H]<sup>+</sup> | 2.83E-08 | 4.28E-06 | 3.31 | Up | 3.31 |
| PC(O-22:0/20:4) | POS | 851.6757 | 852.6835 | 35.52 | [M+H]<sup>+</sup> | 1.04E-04 | 9.67E-04 | 2.70 | Up | 2.70 |
| PC(O-36:5) | POS | 765.5667 | 766.5745 | 21.88 | [M+H]<sup>+</sup> | 3.81E-05 | 5.01E-04 | 2.25 | Up | 2.25 |
| PC(O-38:4) | POS | 795.6114 | 796.6192 | 28.11 | [M+H]<sup>+</sup> | 9.39E-04 | 4.27E-03 | 2.17 | Up | 2.17 |
| PE(O-16:0/22:6) | POS | 749.535 | 750.5428 | 24.18 | [M+H]<sup>+</sup> | 3.28E-03 | 1.09E-02 | 2.40 | Up | 2.40 |
| PS(38:4) | POS | 811.5372 | 812.545 | 22.97 | [M+H]<sup>+</sup> | 1.30E-06 | 5.07E-05 | 3.08 | Up | 3.08 |
| SM(d18:1/16:0) | POS | 702.5663 | 703.5741 | 19.95 | [M+H]<sup>+</sup> | 3.32E-05 | 4.60E-04 | 2.04 | Up | 2.04 |
| GlcCer(d18:1/16:0) | POS | 699.5624 | 700.5702 | 21.05 | [M+H]<sup>+</sup> | 7.30E-05 | 7.59E-04 | 2.61 | Up | 2.61 |
| CE20:4 | POS | 689.6104 | 690.6182 | 49.76 | [M+NH4]<sup>+</sup> | 2.62E-05 | 3.91E-04 | 2.33 | Up | 2.33 |
| TG 53:4 | POS | 885.7803 | 886.7881 | 55.51 | [M+NH4]<sup>+</sup> | 8.84E-07 | 3.96E-05 | 3.92 | Up | 3.92 |
| TG 53:5 | POS | 883.7684 | 884.7762 | 50.97 | [M+NH4]<sup>+</sup> | 1.00E-06 | 4.18E-05 | 4.52 | Up | 4.52 |
| TG 54:5 | POS | 897.7802 | 898.788 | 52.78 | [M+NH4]<sup>+</sup> | 1.74E-04 | 1.37E-03 | 4.14 | Up | 4.14 |
| TG 54:6 | POS | 895.7654 | 896.7732 | 49.98 | [M+NH4]<sup>+</sup> | 2.15E-06 | 7.02E-05 | 2.54 | Up | 2.54 |
| TG 55:4 | POS | 913.8117 | 914.8195 | 62.21 | [M+NH4]<sup>+</sup> | 1.16E-04 | 1.04E-03 | 2.16 | Up | 2.16 |
| TG 55:4 | POS | 913.8109 | 914.8187 | 60.2 | [M+NH4]<sup>+</sup> | 4.81E-08 | 5.54E-06 | 4.29 | Up | 4.29 |
| TG 55:4 | POS | 913.8095 | 914.8173 | 63.62 | [M+NH4]<sup>+</sup> | 2.27E-07 | 1.62E-05 | 6.28 | Up | 6.28 |
| TG 55:6 | POS | 909.7801 | 910.7879 | 49.95 | [M+NH4]<sup>+</sup> | 2.20E-05 | 3.49E-04 | 3.61 | Up | 3.61 |
| TG 55:6 | POS | 909.7812 | 910.789 | 52.7 | [M+NH4]<sup>+</sup> | 5.59E-07 | 2.88E-05 | 6.47 | Up | 6.47 |
| TG 55:7 | POS | 907.7659 | 908.7737 | 47.97 | [M+NH4]<sup>+</sup> | 2.00E-08 | 3.46E-06 | 3.42 | Up | 3.42 |
| TG 56:3 | POS | 929.8412 | 930.849 | 79.24 | [M+NH4]<sup>+</sup> | 8.58E-06 | 1.83E-04 | 3.14 | Up | 3.14 |
| TG 56:4 | POS | 927.827 | 928.8348 | 67.08 | [M+NH4]<sup>+</sup> | 1.53E-05 | 2.73E-04 | 3.61 | Up | 3.61 |
| TG 56:4 | POS | 927.8244 | 928.8322 | 68.81 | [M+NH4]<sup>+</sup> | 2.18E-07 | 1.60E-05 | 5.91 | Up | 5.91 |
| TG 56:5 | POS | 925.8109 | 926.8187 | 59.57 | [M+NH4]<sup>+</sup> | 5.45E-06 | 1.32E-04 | 8.54 | Up | 8.54 |
| TG 54:3 | POS | 901.812 | 902.8198 | 66.65 | [M+NH4]<sup>+</sup> | 4.91E-07 | 2.66E-05 | 3.20 | Up | 3.20 |
| TG 58:7 | POS | 949.8144 | 950.8222 | 56.2 | [M+NH4]<sup>+</sup> | 6.02E-09 | 1.98E-06 | 10.86 | Up | 10.86 |
| TG(18:0_16:0_20:4) | POS | 899.7958 | 900.8036 | 59.22 | [M+NH4]<sup>+</sup> | 3.55E-07 | 2.20E-05 | 5.27 | Up | 5.27 |
| TG(18:1_16:0_22:5) | POS | 923.7959 | 924.8037 | 52.64 | [M+NH4]<sup>+</sup> | 7.28E-09 | 1.98E-06 | 5.84 | Up | 5.84 |
| TG(18:1_16:0_22:6) | POS | 921.7813 | 922.7891 | 50.28 | [M+NH4]<sup>+</sup> | 1.17E-08 | 2.19E-06 | 5.09 | Up | 5.09 |
| TG(18:1_16:0_24:5) | POS | 951.8288 | 952.8366 | 58 | [M+NH4]<sup>+</sup> | 1.70E-07 | 1.33E-05 | 4.23 | Up | 4.23 |
| TG(18:1_18:0_22:5) | POS | 951.8276 | 952.8354 | 59.37 | [M+NH4]<sup>+</sup> | 5.76E-09 | 1.98E-06 | 7.57 | Up | 7.57 |

**Figure 7.**
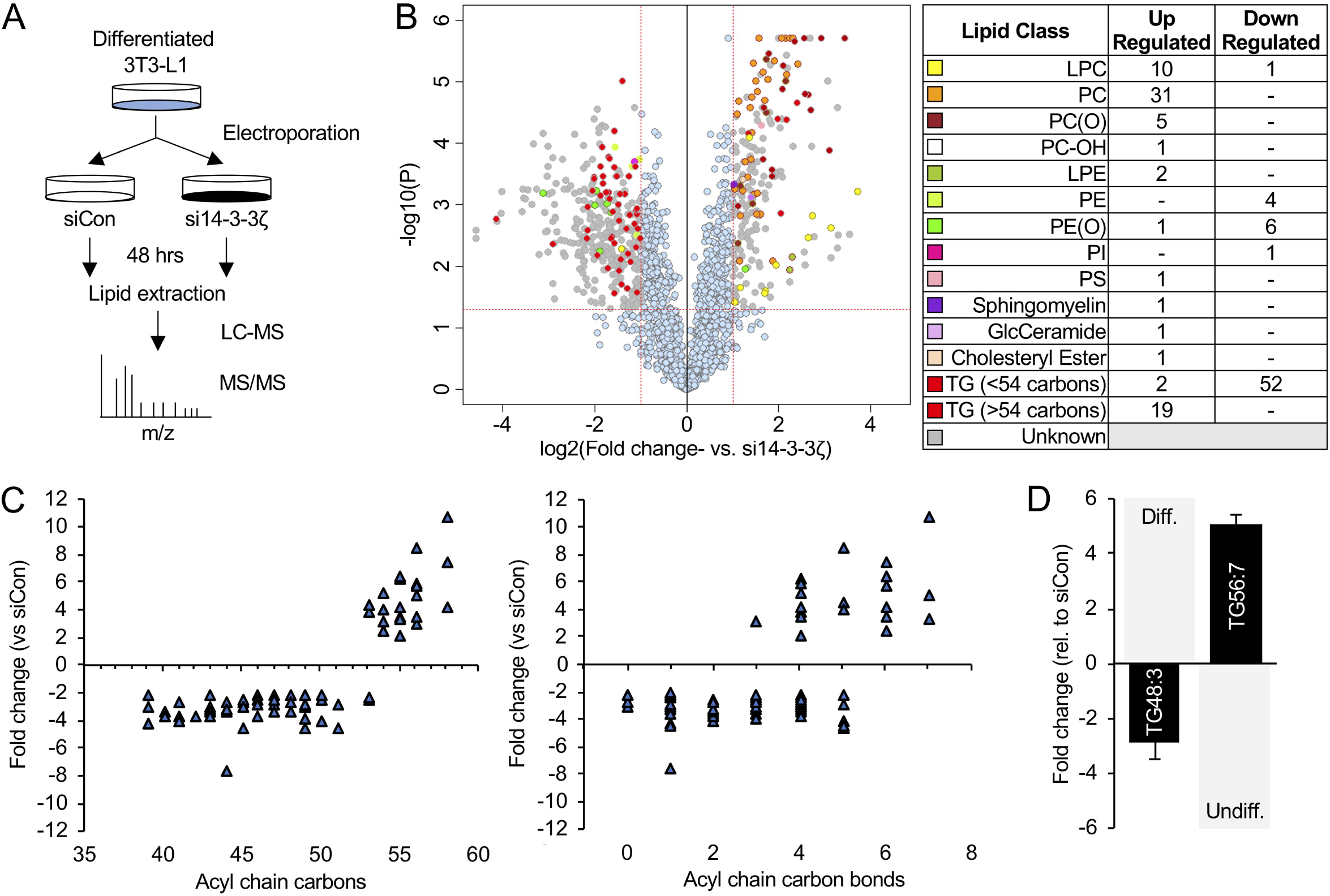
Distinct changes in the lipidome of mature 3T3-L1 cells following 14-3-3ζ knockdown. **(A)** Fully differentiated 3T3-L1 adipocytes were electroporated with control siRNA (siCon) or siRNA against 14-3-3ζ (si14-3-3ζ), and after 48 hours, cells were collected in methanol for lipid extraction and subsequent untargeted LC-MS lipidomics. **(B)** Volcano plot depicting the 2425 features obtained following data processing. The *x* axis corresponds to fold changes (FCs) of MS signal intensity values for all these features in siCon vs si14-3-3ζ cells (in log2) and the *y* axis to *P* values (–log10). Using the selected threshold (corrected *P*-value <0.05 corresponding to a FDR<5% and FC>2 or <0,5), 549 features were found to significantly discriminate siCon vs si14-3-3ζ cells, of which 139 unique lipids from various (sub)classes (shown using color symbols) were identified using an in-house reference database. The in-set table lists the number of lipid species that were found to be up- or down-regulated according to the different lipid (sub)classes following 14-3-3ζ depletion. **(C)** Triglycerides that were significantly up- or down-regulated after 14-3-3ζ knockdown were categorized by acyl carbon chain length or by the degree of saturation. **(D)** Triglycerides species that are normally enriched (TG 48:n) or depleted (TG 56:n) in differentiated 3T3-L1 cells (gray columns) were found to be significantly affected by 14-3-3ζ depletion (n=5 per group). Values represent fold-change ± SE.

It has been documented that following the differentiation of 3T3-L1 adipocytes, specific triglyceride species, like those with acyl carbon chain lengths ≥48 carbon lengths and those containing mono- and polyunsaturated species like TG 48:1, 48:2, and 48:3, are increased, whereas certain triglyceride species with 56 acyl carbons and 7 unsaturated bonds are decreased (33). Of note, knockdown of 14-3-3ζ in mature 3T3-L1 adipocytes led to significant decreases in TG 48:n species and the converse enrichment of TG 56:n species, both of which are associated with a pre-adipocyte lipidome signature (Figure 7D; Table 1) (33). When taken together, these findings further demonstrate that 14-3-3ζ depletion promotes the acquisition of an immature phenotype.

## Discussion

As 14-3-3ζ and its related isoforms are implicated in diverse processes, the initial intent of the present study was to examine whether 14-3-3ζ could influence metabolic processes in mature adipocytes. This would build on the work of others and our own studies that identified essential roles of 14-3-3ζ and its related isoforms in adipogenesis, glucose uptake, and oxidative metabolism (7, 8, 12, 34, 39, 43-45). We focused primarily on lipolysis, as the lipolytic enzyme HSL is phosphorylated by PKA to create 14-3-3 protein binding sites, and found that reductions in 14-3-3ζ expression was sufficient to attenuate lipolysis (1, 36). Unexpectedly, we found that 14-3-3ζ was not necessary for β3-adrenergic receptor-mediated lipolysis; instead, reducing 14-3-3ζ levels promoted the acquisition of an immature adipocyte phenotype that was associated with significant decreases in the expression of lipolytic, lipogenic, and functional adipocyte genes. These adipocytes displayed significantly impaired responses to lipolytic stimuli, as a result of the observed immature phenotype. Collectively, this study highlights novel roles of 14-3-3ζ in the regulation of adipocyte maturity.

Most striking was the observation that 14-3-3ζ could influence adipocyte maturity *in vitro* and *in vivo.* We found that reducing 14-3-3ζ expression in fully mature adipocytes significantly reduced PPARγ mRNA and protein levels and genes associated with mature adipocytes. When taken with our previous work where we described essential functions of 14-3-3ζ in adipogenesis and adipocyte differentiation (34), it appears that14-3-3ζ represents a critical factor in regulating the cellular fate of pre- and mature-adipocytes, as well as the maintenance of adipocyte identity and maturity. Adipocyte dedifferentiation has been documented in pregnancy, tissue repair, and tumorigenic transformations of liposarcomas (4, 10, 32), and dedifferentiation of visceral and subcutaneous human adipocytes, as noted by decreased mRNA expression of mature adipocyte gene markers including *PPARG2, C/EBPA, LPL*, and *ADIPOQ*, has also been documented (37). The cellular mechanisms underlying adipocyte dedifferentiation can be mediated by Notch receptor activation or canonical and non-canonical WNT signaling pathways (5, 22, 52). For example, in 3T3-L1 cells, WNT3A-associated dedifferentiation occurs by increased levels of β-catenin, which results in the induction of specific genes leading to a loss of adipocyte identify (22). As 14-3-3ζ has been shown to be a negative regulator of β-catenin localization and expression, reducing 14-3-3ζ abundance may promote dedifferentiation through a similar mechanism (31). Further work is required to determine whether immature 14-3-3ζ-expressing adipocytes can re-differentiate into mature adipocytes or transdifferentiate into other cell types, such as macrophages, endothelial cells, or osteoblasts (22, 27, 47, 48).

In addition to affecting adipocyte maturity, depletion of 14-3-3ζ was also associated with marked decreases in triglyceride content and genes associated with lipid synthesis. In differentiating 3T3-L1 cells, distinct changes in the lipidome signature have been reported, whereby certain lipid species, such as triglycerides of varying acyl carbon lengths, are increased or decreased during differentiation (33). Our unbiased lipidomic analysis revealed over 139 unique lipid species, predominantly triglycerides, whose abundance were significantly changed following 14-3-3ζ knockdown. Reducing 14-3-3ζ expression in mature adipocytes decreased levels of triglyceride species with shorter chain (≤ 54) lengths and low number of unsaturations and increased species with longer chain (>54) lengths and high number of unsaturations (33). Most notable was the finding that 14-3-3ζ depletion promoted the acquisition of a specific lipid signature where triglycerides species associated with fully differentiated 3T3-L1 adipocytes were decreased, while increased abundance of those found in pre-adipocytes was observed (33). The exact mechanisms by which 14-3-3ζ knockdown in 3T3-L1 cells reduces intracellular lipids is not known, but it is possible that autophagy, which we have previously shown to be up-regulated by 14-3-3ζ depletion, could be a potential contributor (15, 34). Alternatively, dedifferentiation of adipocytes is also associated with a rapid expulsion of lipid droplets from cells via a yet undefined mechanism (11).

Within a given adipose tissue depot, it is now appreciated that heterogeneous populations of adipocytes exist, and these adipocytes can arise from different progenitors pools, such as those expressing DPP4, WT1, and PDFRα (24, 30, 40, 46, 50). In the present study, we found that 14-3-3ζ is heterogeneously expressed within a sub-set of adipocytes in gWAT of male mice. The loss of 14-3-3ζ in these cells could account for the small, but significant, reductions in the expression of adipocyte maturity, lipolytic, and lipogenic genes in gWAT. The heterogenous expression of 14-3-3ζ presents a limitation in purifying sufficient numbers of 14-3-3ζ-expressing adipocytes from gWAT to directly assess the role of 14-3-3ζ, and to circumvent this issue, appropriate lineage-tracing fluorescent reporter mice are clearly needed, as such tools would be invaluable to purifying 14-3-3ζ-expressing adipocytes (30). In the present study, 3T3-L1 cells were primarily used to understand how 14-3-3ζ influences adipocyte maturity, as they recapitulate many aspects of primary murine adipocytes (34, 42). We acknowledge the limitation that they are not bonafide adipocyte pre-cursor cells that are found within the stromal vascular fraction of gonadal adipose tissues, but they represented a homoegenous cell population that allowed the direct examination how 14-3-3ζ influences maturity.

Additional studies are required to understand whether 14-3-3ζ can have additional roles in mature adipocytes or in the context of glucose homeostasis. For example, we previously reported that systemic 14-3-3ζ knockout mice displayed reduced adiposity, whereas transgenic mice overexpressing 14-3-3ζ gained more weight on a high-fat diet. When taken together, this suggests that 14-3-3ζ could influence high-fat diet-associated adipose tissue expansion (34). It has been established that decreased PPARγ expression in mouse adipocytes impairs WAT and BAT formation, as well as diet-induced obesity (29, 51), and with our observaion that adipocyte-specific deletion of 14-3-3ζ reduced PPARγ2 levels in gWAT (Figure 5D), deletion of 14-3-3ζ may confer protection to diet-induced obesity. However, any effects on weight gain could be due to the inability of immature adipocytes to store lipids in times of nutrient excess. Additionally, the regulatory roles of 14-3-3 proteins on AS160/TBC1D4-associated glucose uptake have been well-documented *in vitro* and *in vivo* (9, 45). Based on our previous finding that systemic 14-3-3ζ knockout mice had decreased insulin sensitivity (34), we had hypothesized that deletion of 14-3-3ζ in adipocytes would cause adipocyte-specific impairments in GLUT4 translocation, thereby affecting glucose uptake. In the present study, no differences in glucose tolerance or insulin sensitivity in adi14-3-3ζKO mice were observed, which suggests that the decrease in insulin sensitivity observed in systemic 14-3-3ζKO mice could be specific to skeletal muscle, which is the primary site of glucose disposal (13). Thus, additional studies are required to assess the roles of 14-3-3ζ in skeletal muscle.

Collectively, results from our *in vivo* and *in vitro* models demonstrate that 14-3-3ζ is required for the maintenance of adipocyte maturity and this results in an impairment in adipocyte function. To date, the functional roles of 14-3-3ζ in pre-adipocytes and mature adipocytes are not fully known, but when taken with our previous work, this study highlights the complexity of 14-3-3ζ. Additional studies are needed to further characterize the differences between adipocytes that do and do not express 14-3-3ζ, and increasing our understanding of how 14-3-3ζ may regulate adipocyte function may lead to the development of novel approaches to treat lipid abnormalities stemming from obesity.

## Abbreviations

ATGL: adipose triacylglycerol lipase
HSL: hormone-sensitive lipase
MAGL: monoacylglycerol lipase
PKA: protein kinase A
cAMP: cyclic adenosine monophosphate

## Acknowledgments

The authors would like to thank Mina Sadeghi and Dr. Yves Mugabo (CRCHUM) for their technical assistance with various experiments. The authors would also like to thank Dr. Amparo Acker-Palmer (Institute for Cell Biology and Neuroscience, Goethe University) for providing breeding pairs of TAP-14-3-3ζ transgenic mice. The authors would also like to thank the CRCHUM Histological core and the Transgenesis and Animal Modeling Core for assistance with histology and the rederivation of *Ywhaz*^flox/flox^ mice, respectively.

## Authors contributions

A.O. designed the studies, carried out the research, interpreted the results, and wrote the manuscript. K.D. performed experiments and edited the manuscript. IRF performed experiments and analyzed data. C.D. contributed to study design and edited the manuscript. G.E.L. conceived the concept, designed and performed studies, interpreted the results, and wrote and revised the manuscript. G.E.L is the guarantor of this work.

## Supplemental Figure legends

**Figure S1- Deletion of 14-3-3ζ in adipose tissue of female mice does not promote metabolic dysfunction. (A)** Levels of *Ywhaz* mRNA were measured by quantitative PCR in gonadal (gWAT) and inguinal (iWAT) adipose tissue from female adi14-3-3ζKO mice (n=8 per genotype). **(B)** Body weights of female adi14-3-3ζKO mice following i.p injections of tamoxifen (50 mg/kg b.w) were measured weekly (n=8 per genotype). **(C**,**D)** Intraperitoneal glucose (2 g/kg b.w.) (D) and insulin (0.5 U/kg b.w.) (E) tolerance tests on female adi14-3-3ζKO mice fasted for 6 and 4 hours, respectively, at 12 weeks of age (n=8 per genotype). **(E**,**F)** Plasma glycerol and FFA levels in female adi14-3-3ζKO (E) and TAP (F) mice after injection with CL-316,243 (CL, 1 mg/kg b.w.) or isoproterenol (ISO, 1 mg/kg b.w.) following an overnight fast (adi14-3-3ζKO: n=8 WT, n=8 KO; TAP: n=9 WT, n=12 TAP). All values represent mean ± SE.

**Figure S2- Over-expression of 14-3-3ζ does not affect lipolysis in male mice. (A)** After an overnight fast, body weights from male wild-type (WT) and TAP-14-3-3ζ over-expressing transgenic mice (TAP) were obtained (n=6 WT, n=9 TAP mice). **(B)** Following an overnight fast, plasma glycerol and FFA levels in male wild-type (WT) and TAP-14-3-3ζ over-expressing transgenic mice (TAP) were measured after an intraperitoneal injection with isoproterenol (ISO; 10 mg/kg b.w.) (n=6 WT, n=9 TAP mice). **(C)** Mature 3T3-L1 adipocytes were transfected with plasmids containing GFP (2 μg) or 14-3-3ζ IRES-GFP (2 μg), and over-expression was confirmed by immunoblotting (n=4 per group). **(D)** Free fatty acid (FFA) release into the supernatant after a 2-hour incubation with isoproterenol (ISO, 1 μM.) was measured from GFP or 14-3-3ζ IRES-GFP 3T3-L1 adipocytes (n=4 per group; two-way ANOVA, followed by Bonferroni t-tests; *: p<0.05 when compared to siCon+ISO). All values represent mean ± SE.

---

**Supplemental Table 1.**
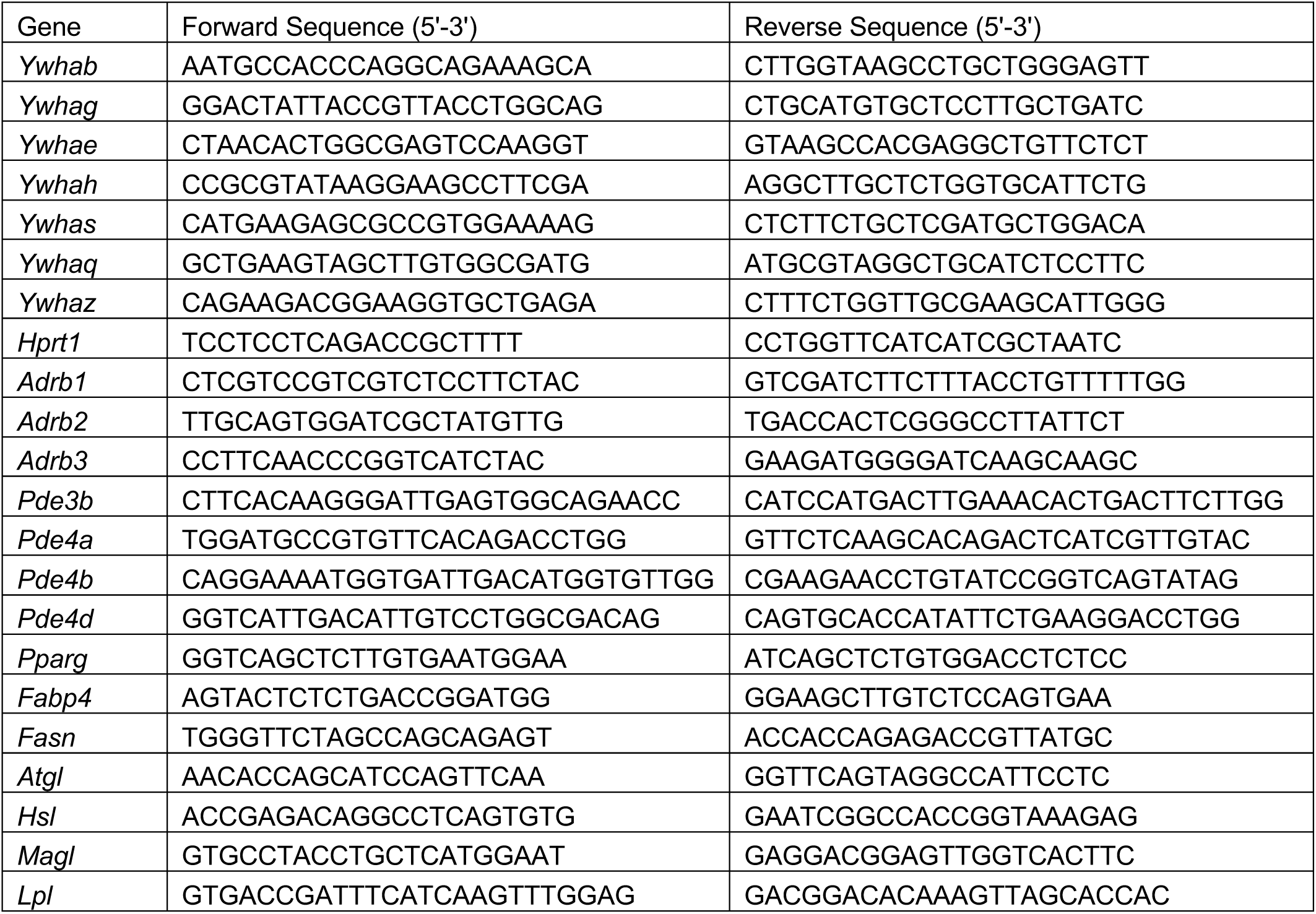
Primer sequences used for qPCR.

**Supplemental table 2:**
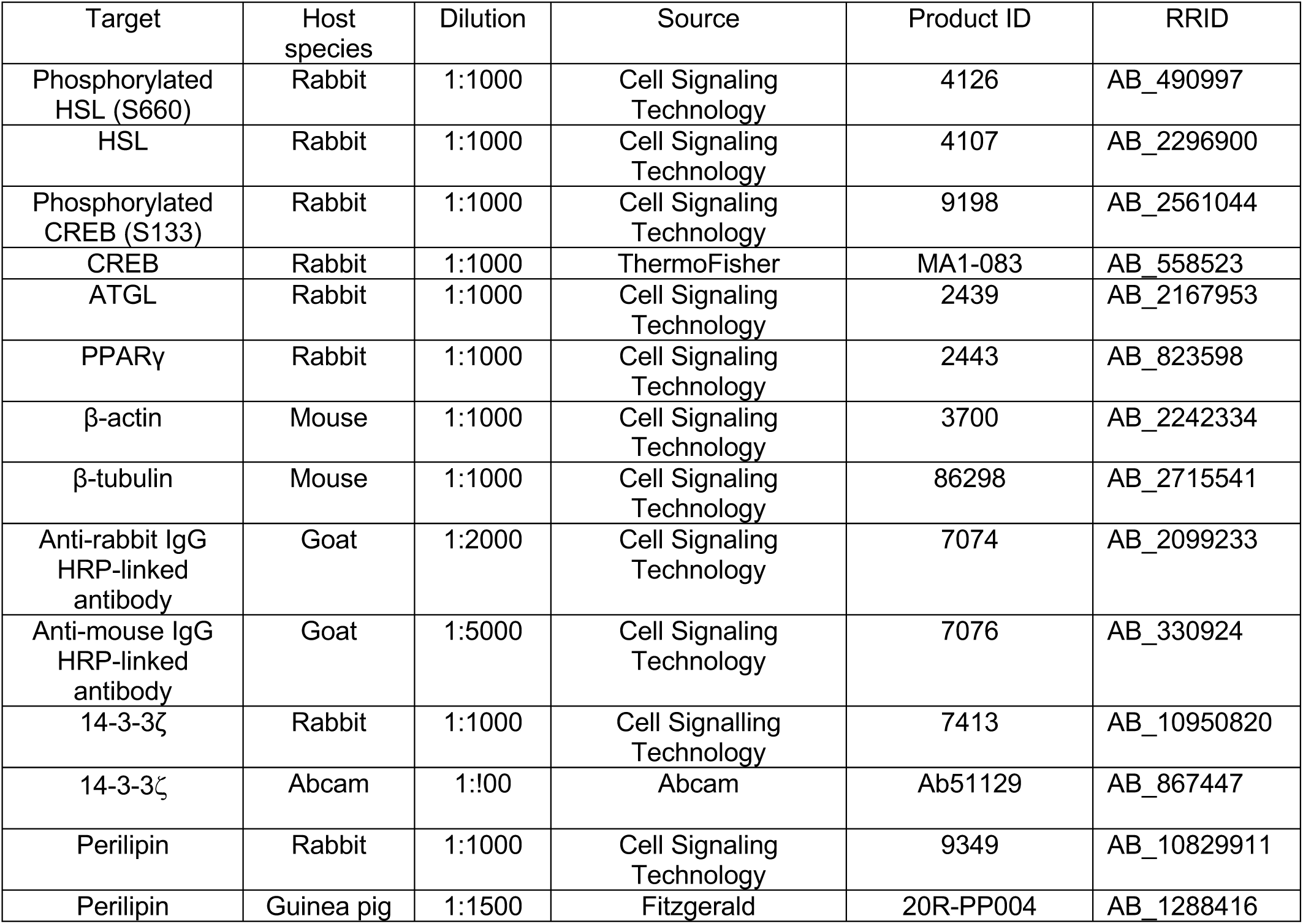
Antibodies used.

