## Supplementary figures and images for "Reducing 14-3-3ζ expression influences adipocyte maturity and impairs function" (from another version of this preprint: version 1, 2020-02-25)

### Supplemental Figures

#### Supplemental Figure 1

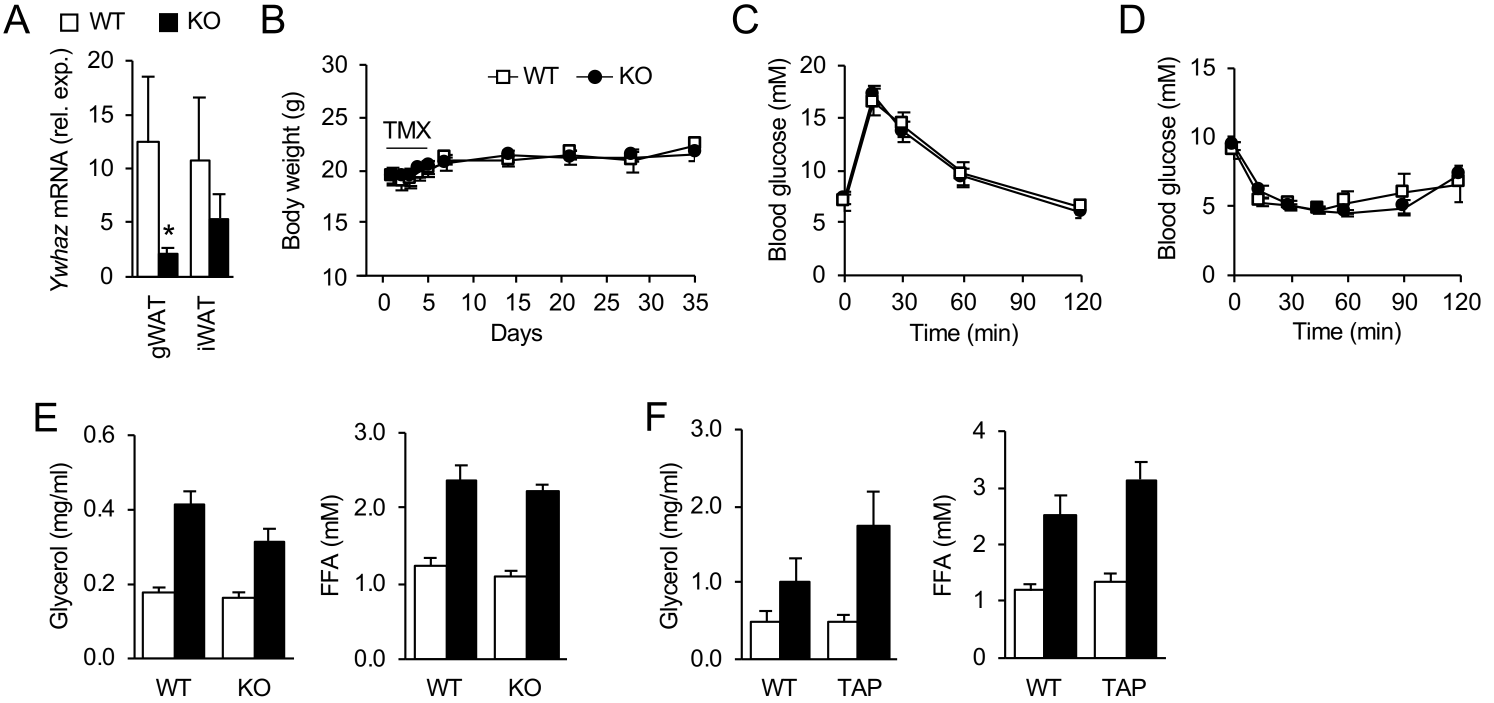

#### Supplemental Figure 2

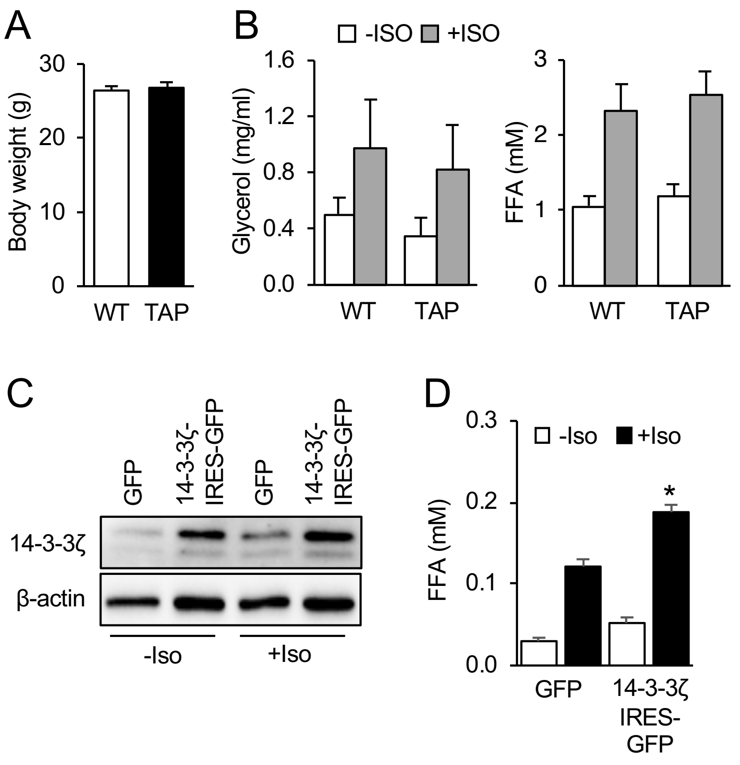
